# Src promotes tumor cell invasion by hijacking the translation machinery

**DOI:** 10.1101/2024.08.01.606119

**Authors:** Benjamin Bonnard, Anouk Chatefau, Cyril Dourthe, Sylvaine Di Tommaso, Jean-William Dupuy, Isabelle Mahouche, Jacobo Solorzano, Morgane Le Bras, Anne-Aurélie Raymond, Violaine Moreau, Arnaud Jabouille, Anne Blangy, Yvan Martineau, Frédéric Saltel

**Affiliations:** Bordeaux Institute of Oncology, BRIC U1312, INSERM, Université de Bordeaux, Bordeaux, France. Labeled LNCC; Oncoprot Platform, TBM-Core US 005, Bordeaux, France; Centre de Recherche en Cancérologie de Toulouse (CRCT), INSERM U1037, Université Toulouse III Paul Sabatier, ERL5294 CNRS, Toulouse, France; Montpellier Cell Biology research center, CRBM, Univ Montpellier, CNRS, Montpellier, France

**Keywords:** Cancer, invasion, invadosomes, Src, translation initiation, eIF3

## Abstract

The Src oncogene controls cancer cell invasiveness by promoting invadosome formation and extracellular matrix degradation (ECM). Invadosomes are enriched in the eukaryotic translation initiation factor 3 (eIF3) complex associated with a local mRNA translation activity mandatory for their maintenance. Here, we show that Src regulates mRNA translation by controlling the expression of eIF3 subunits. Among them, eIF3h/e/d are essential for invadosome formation and ECM degradation. We demonstrate that Src controls the canonical mTOR/eIF4E and the non-canonical eIF3d cap-dependent translation initiation pathways. We show that both pathways are necessary for invadosome formation and their ECM degradation function. Finally, we highlighted a correlation between *Src* and *eIF3h/e/d* overexpression, which is associated with poor prognosis in hepatocellular carcinoma (HCC) patients and controls the ECM degradation and invasive properties of HCC cells. These findings identify Src as a major regulator of translation initiation pathways, which leads to invadosome formation, ECM degradation and tumor cell invasion.

Graphical abstract

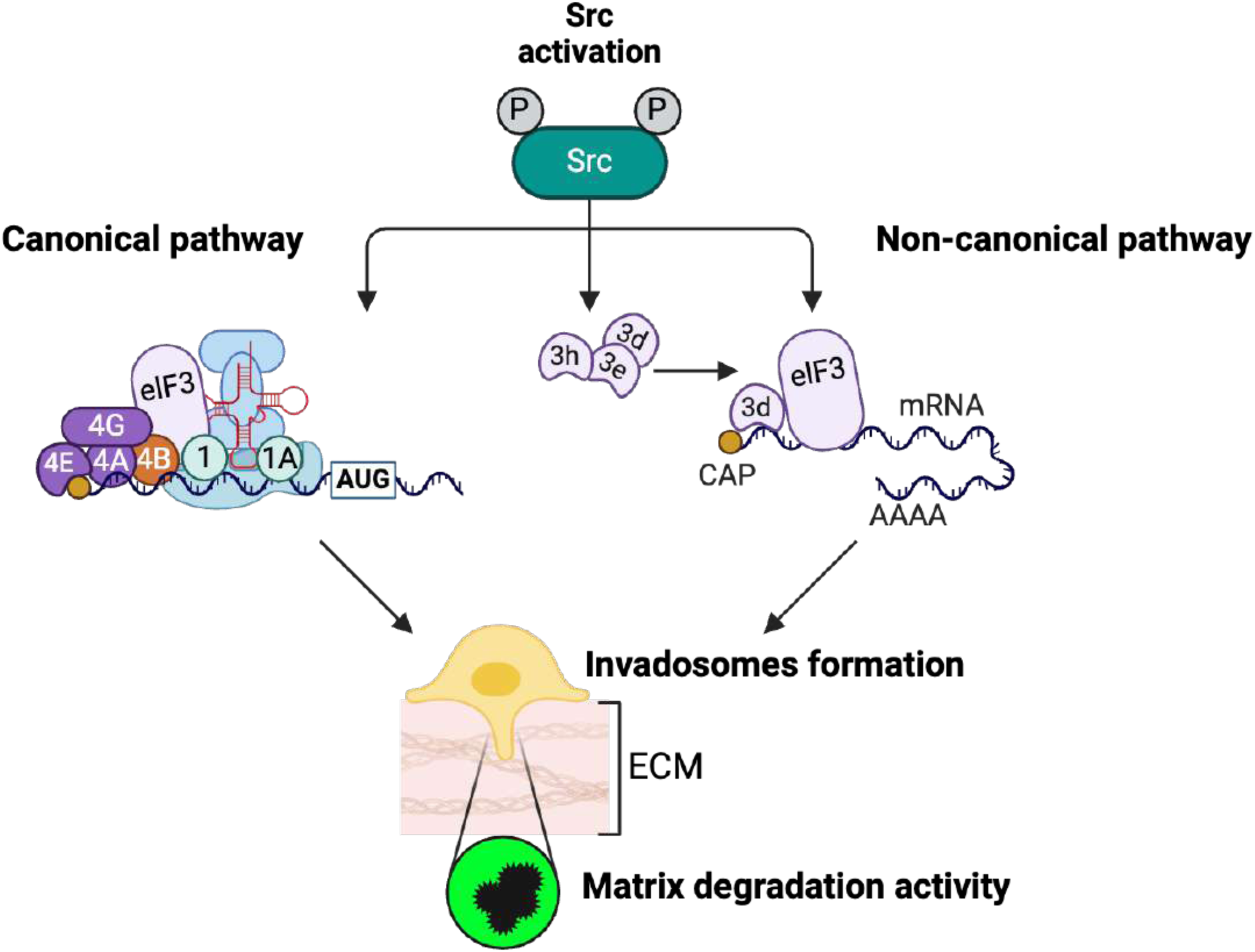

## Introduction

The degradation of the extracellular matrix (ECM) is a determinant step for cancer cell migration and invasion leading to tumor progression and metastasis development^1^. Invadosomes are dynamic F-actin based structures mainly involved in ECM remodeling in normal and cancer cells^2,3^. As observed for cell migration^4,5^, cell protrusions require the re-localization of messenger RNA (mRNA) and translation^6,7^. Previously, we demonstrated the local enrichment of the translational machinery (including ribosomes, β-actin mRNA) in invadosomes from Rous sarcoma virus transformed NIH-3T3 cells (NIH-3T3 Src), associated with a local mRNA translation activity^8^. Furthermore, we found that several translation initiation factors were enriched in invadosomes as compared to the whole cell, including 8 out of the 13 subunits of the eukaryotic translation initiation factor 3 (eIF3) complex, suggesting a determining role of the eIF3 complex in invadosome formation^8^. The aberrant expression of individual eIF3 subunit proteins and mRNAs was reported in several cancers including liver, lung, breast and colorectal cancers^9^. The depletion of individual eIF3 subunits (a/b/c/e/g/i) was found to reduce global translation and cell proliferation^10^. Some studies have started to described a potential invasive role of eIF3 subunits in cancer cells. Thus, it has been shown that eIF3h-depleted HCC LM3 cell line had lower invasive capacity^11^, and eIF3d promotes breast cancer invasion and metastasis through its non-canonical cap-binding activity in association with eIF4g2, a distinct variant of eIF4g^12^. Moreover, the overexpression of individual eIF3 subunits (a/b/c/h/i) was also shown to enhance NIH-3T3 cell proliferation^13^. Nevertheless, the potential role of eIF3 complex in invadosome formation and the exact molecular mechanism involved remain to be elucidated.

The eIF3 complex plays a major role in translation initiation, which involves the recruitment of ribosomes to the mRNA and requires the coordinated interaction between the eIF4f (composed by cap-binding protein eIF4e, the scaffold protein eIF4g and the RNA-helicase eIF4a) and the eIF3 complexes. The eIF3 complex promotes the assembly of the 43S pre-initiation complex, composed of the 40S ribosomal subunit and numerous others eIFs (including eIF1, eIF1a and eIF2), strengthening mRNA binding to the eIF4f complex and scanning for the start codon^9^. The initiation of cap-dependent mRNA translation is controlled by the PI3K/mTOR pathway^14^, which can be activated by the tyrosine kinase Src^15^. Src kinase activity was shown to drive the translation of polycomb complex components in breast cancer tumors, leading to their overexpression^16^. Nevertheless, the direct link between Src and the translation machinery remains unclear.

In this study, we demonstrate that Src activity regulates protein synthesis through activation of both the canonical and the non-canonical mRNA translation pathways and that it controls the expression of specific eIF3 complex subunits. Furthermore, we identify individual eIF3 subunits that are essential for invadosome formation and ECM degradation. We performed next-generation sequencing on laser captured invadosomes and identified eIF3-dependent target mRNAs involved in invadosome formation: the mRNAs of the RNA binding protein IGF2BP2 (insulin like growth factor 2 mRNA-binding protein 2) and of transketolase (TKT), an enzyme involved in pentose-phosphate pathway. Both IGF2BP2 and TKT proteins localize into invadosomes and their depletion prevents invadosome formation. In addition, we highlight that *Src* and *eIF3* are enriched in tumor section from hepatocellular carcinoma (HCC) patient biopsies and we demonstrate that Src and eIF3 participate in HCC cell invasiveness. Together, our study identifies that the Src/mRNA translation initiation pathway is essential for to invadosome formation and cancer cell invasion.

## Results

### Src kinase activity regulates protein synthesis and specific eIF3 subunit expression

Src-induced invadosomes are a hotspot for translation with the enrichment of β-actin mRNA into these invasive structures^8^. To confirm the colocalization of both β-actin mRNA and protein in invadosomes of in NIH-3T3 Src cells, we used the Halo-actin translation reporter previously used in neurons^17^. By co-expression of the Halo-Tag β-actin reporter and the MS2 RNA stem-loop binding protein stdMCP-stdGFP, we observed that both Halo-Tag β-actin reporter mRNA and protein colocalized at invadosomes (Figure 1A). This supports our hypothesis that local translation of mRNAs occurs in invadosomes. Since the pioneer studies^18,19^ showing that the kinase Src is a key driver for most of invadosomes^20^, Src signaling has been described to be involved in several cellular functions including proliferation, angiogenesis, cytoskeleton organization^21^, with a potential role in protein synthesis^15^. In fact, we found that NIH-3T3 Src cells, which express the constitutively active form of Src, have an overall translation level higher than classical NIH-3T3 WT cells, as assessed by SUrface SEnsing of translation (SUnSET) assays (Figure 1B). To confirm the involvement of Src activity is in translation, we used a pharmacological approach with Src kinase inhibitors (PP2 or SU6656), as well as a targeted approach using siRNAs in NIH-3T3 Src cells. Short-term (1 hour) pharmacological treatment with Src inhibitors or Src depletion with siRNAs reduced Src activity (Figure 1C, S1A and S1D) and resulted in decreased invadosome formation (Figure 1D and S1B) and protein synthesis levels (Figure 1E, S1C and S1E). Interestingly, Src reactivation after drug elimination restored both invadosome formation and translation (Figure 1C-E and S1A-1D). We also found that Src activation level is positively correlated with translation level in NIH-3T3 Src cells (Figure 1E, right panel). Using polysome profiling, we confirmed that Src activation regulated the association between mRNAs and ribosomes: PP2 treatment caused loss of heavy polysome and a corresponding increase of 60S and 80S fractions (Figure S1F). This suggests that analyzing mRNA abundance was not sufficient to measure the fully measure the impact of Src inhibition. Therefore, we looked at proteome changes following Src expression to further decipher modified cell processes.

**Figure 1:**
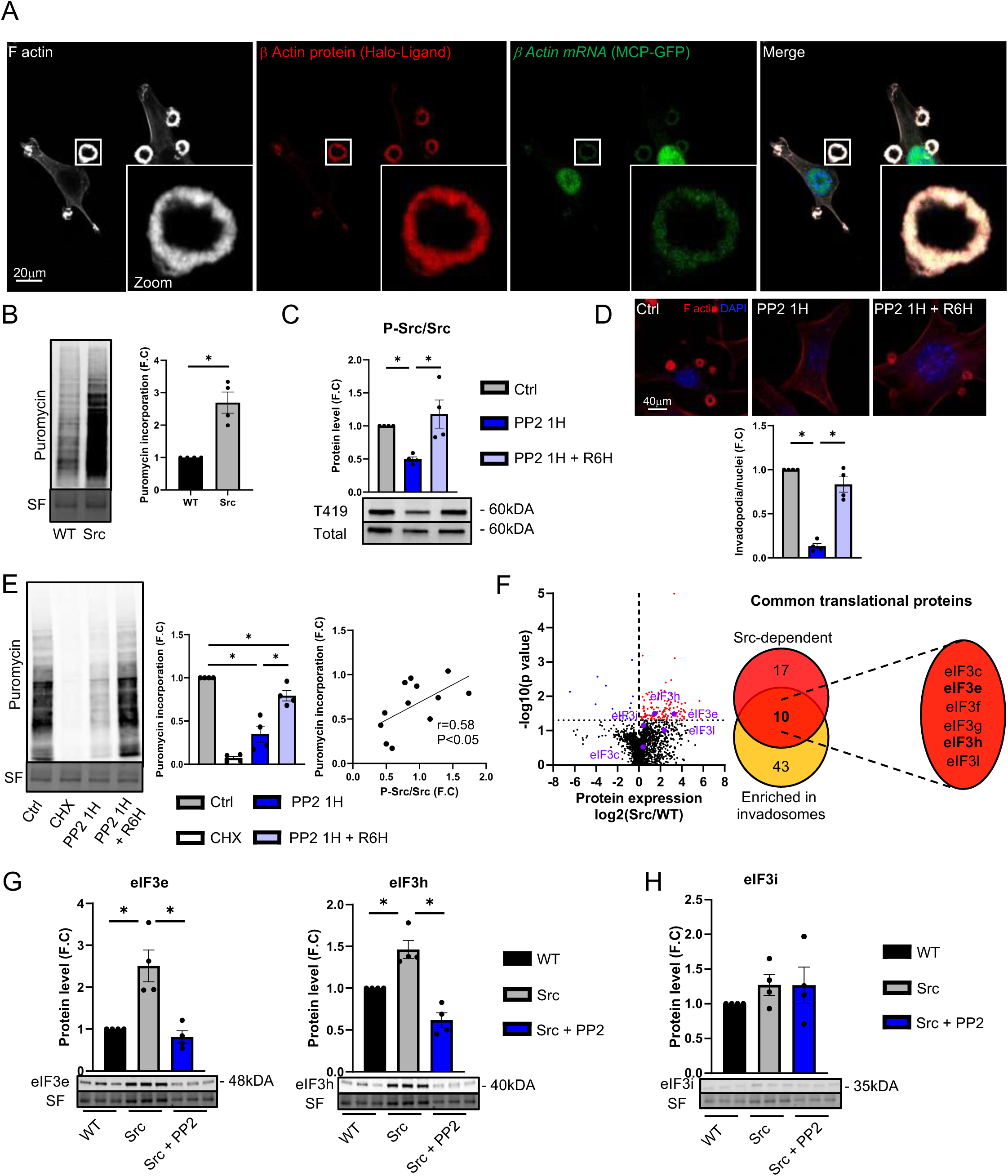
Src kinase activity regulates protein synthesis and specific eIF3 subunit expression. (A) NIH-3T3 Src cells were transfected with HaloTag β−actin reporter associated with stdMCP-stdGFP. While β−actin mRNA was detected by GFP staining (green), the incubation with JF646 Halo-ligand revealed the localization β−actin protein (red) (scale bar: 20μm). (B) The Surface Sensing of Translation (SUnSET) assay was conducted on transfected NIH-3T3 WT (WT) and NIH-3T3 Src (Src) cells. The bar graph depicts the quantification of puromycin incorporation. (C-E) Src cells underwent treatment with Src inhibitor for 1 hour (PP2 1H) followed by a recovery period in fresh medium (PP2 1H + R6H). DMSO treatment served as the control group (Ctrl). (C) Src activity was evaluated by western blot. (D) The right panel displays representative images of invadosomes stained with phalloidin, with the accompanying bar graph indicating the number of invadosomes per nucleus. Scale bar: 40μm. Error bars (mean ± SEM, *n*=20 fields, four independent experiments, *P<0.05, paired t-test was used for statistical analysis). (E) The SUnSET assay was performed on treated or untreated Src cells. As a negative control, Src cells were incubated with cycloheximide (CHX). The middle panel illustrates the quantification of puromycin incorporation, while the right panel displays the Pearson correlation between Src phosphorylation and puromycin incorporation in Src cells challenged or not by PP2 1H or PP2 1H + R6H. (F) The volcano plot highlighting downregulated (blue) and upregulated (red) proteins in Src cells compared to WT cells, evaluated by mass spectrometry. Protein expression of eIF3c/e/h/i/l subunits is indicated in purple. The Venn diagram illustrates the overlap between deregulated translational proteins resulting from Src activation and those enriched into invadosomes compared to whole cells. (G-H) Protein expression levels of eIF3h, eIF3e and eIF3i in WT and Src cells treated or untreated with PP2 or for 24 hours. Bar graphs for western blot quantification were presented with error bars (mean ± SEM, four independent experiments, *P<0.05, paired t-test was used for statistical analysis).

To further decipher Src-dependent proteome, we performed a comparative proteomic analysis by mass spectrometry of NIH-3T3 WT and NIH-3T3 Src cells, treated or not with PP2. We found 135 proteins were upregulated in NIH-3T3 Src as compared to NIH-3T3 WT cells, while only 15 proteins were downregulated (Figure 1F, left panel). Conversely, PP2 treatment in NIH-3T3 Src cells downregulated 106 proteins and upregulated 38 (Figure S1G). In order to better understand to role of Src activity in the regulation of invadosome proteome, we compared the Src-dependent proteome, i.e. proteins upregulated in NIH-3T3 Src as compared to NIH-3T3 WT cells (Figure 1F, left panel) or downregulated by Src inhibition (Figure S1G) to the proteins we found enriched in the proteome of invadosomes^8^. By gene ontology analysis, we found that 27 proteins regulating mRNA translation were Src-dependent, among which 10 were enriched in invadosomes (Figure 1F, right panel; Table S1), including six subunits of the eukaryotic initiation factor 3 (eIF3) complex: eIF3c, e, f, g, h and l. The largest translation initiation factor eIF3 has a determinant role for protein synthesis and we confirmed by immunoblot that Src activity regulates the expression of certain eIF3 subunits. Indeed, we found that eIF3e and eIF3h are expressed at higher levels in NIH-3T3 Src cells as compared to NIH-3T3 WT cells (Figure 1G and S1H). Moreover, the inhibition of Src activity reduced expression of eIF3c, e, h and l (Figure 1G, S1I and S1J). Conversely, we found that Src activity did not regulate the expression of eIF3i (Figure 1H and S1K), suggesting that Src controls the expression of specific subunits of eIF3 in NIH-3T3 Src cells.

Taken together these results demonstrate that Src is a determining driver of protein synthesis, by promoting the expression of a subset of eIF3 subunits.

### Key role of Src-regulated eIF3 subunits in invadosome formation and ECM degradation

The eIF3 complex is composed by 13 different subunits and 8 eIF3 subunits were found enriched in invadosomes^8^, as depicted in Figure 2A. To understand the role of Src-dependent eIF3 subunits (eIF3h and eIF3e) in invadosome formation and matrix degradation activity, we assessed the sub-cellular localization of eIF3h and eIF3e during invadosome formation, by expressing eIF3h-GFP or eIF3e-GFP in NIH-3T3 Src cells. We found that eIF3h and eIF3e concentrated in the center and the periphery of invadosome rosettes (Figure 2B and S2A), similarly to the endoplasmic reticulum as described previously^8^. In addition, eIF3h co-localized with invadosome during all its development steps, from initiation to collapse (Figure 2C and S2A; Video 1 and 2). This suggested a role of eIF3 subunits for invadosome formation.

**Figure 2:**
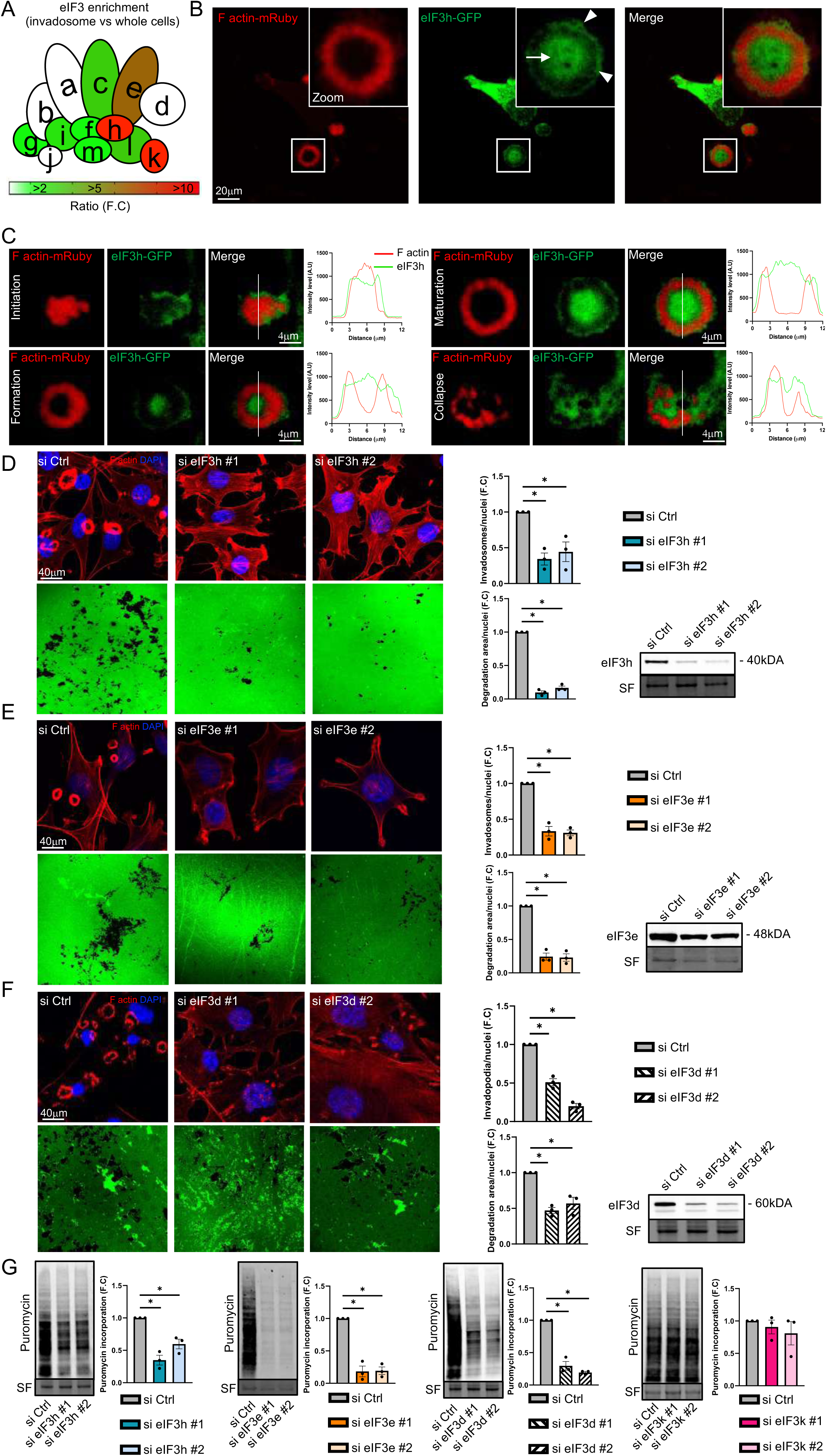
Key role of Src-regulated eIF3 subunits in invadosome formation and ECM degradation. (A) Schematic representation of eucaryotic translation imitation factor 3 (eIF3) complex showing the individual enrichment of eIF3 subunit into invadosomes vs whole cells from Ezzoukhry. Z *et al*^1^. Colored bar represents enrichment ratio (invadosome vs whole cells) from <2 (white) to >10 (red). (B-C) Representative images from time-lapse video microscopy of lifeact-mRuby (red)-expressing NIH-3T3 Src cells transfected with eIF3h-GFP (green) in matured invadosome (b, scale bar: 20μm) and during invadosome formation (c, scale bar: 4μm). (B) Arrow and heads of arrow indicate the sub-localization of eIF3h in center and in periphery respectively of invadosome. (C) Bar graphs show F-actin and eIF3h intensity levels as a function of defined distance axis (indicated by white line). (d-g) NIH-3T3 Src cells were transfected with siRNA control (si Ctrl) or with two independent siRNAs targeting eIF3h (D) or eIF3e (E) or eIF3d (F). (D-F) Transfected cells were seeded on a fluorescent gelatin matrix. The upper panel shows representative images of invadosomes stained with phalloidin; the bar graph shows the number of invadosomes per nuclei. The lower panel shows representative images of degradation area (black); the bar graph shows the quantification of matrix degradation area per nuclei after 24 hours. Scale bar: 40μm. Error bars (mean ± SEM, *n*=20 fields, three independent experiments, *P<0.05, paired t-test was used for statistical analysis). (G) SUnSET assay performed on transfected NIH-3T3 Src cells with si Ctrl or siRNAs targeting eIF3h or eIF3e or eIF3d. Bar graphs show the quantification of puromycin incorporation. Error bars (mean ± SEM, three independent experiments, *P<0.05, paired t-test was used for statistical analysis).

To determine the role of eIF3h and eIF3e in invadosome formation and ECM degradation activity, we individually depleted their expression by using small interfering RNA (siRNA) in NIH-3T3 Src cells. We found that individual the depletion of either eIF3h or eIF3e reduced invadosome formation, ECM degradation and cell invasiveness (Figure 2D, 2E, S2B and S2C). Because eIF3h is required for the assembly of eIF3k/l/e/d subunits and eIF3e for the assembly of eIF3k/l/d on the eIF3 complex^22^, we assessed whether eIF3h and eIF3e depletion could affect the other eIF3 subunits expression. We found that eIF3h depletion reduced the levels of eIF3e, d and k, whereas the depletion of eIF3e reduced expression of eIF3d and k without affecting that of core eIF3 subunits such as eIF3a, b and c (Figure S2D and S2E). These results suggest that eIF3 subunits eIF3d and eIF3k, which are regulated by eIF3h and eIF3e, could also participate in invadosome formation.

By individual depletion of eIF3d or eIF3k in NIH-3T3 Src cells as above, we found that the depletion of eIF3d reduced invadosome formation, ECM degradation and cell invasiveness, while the depletion of eIF3k had no effect (Figure 2F, S2F and S2G). We observed that the individually depletion of eIF3d or eIF3k did not affect the expression of eIF3h and eIF3e (Figure S2H and S2I). The depletion of eIF3d led to higher levels of eIF3c but did not affect the other core eIF3 subunits, and the depletion of eIF3k did not change the level of any of the core eIF3 subunits (Figure S2H and S2I). These results demonstrate that eIF3h, eIF3e and eIF3d have a crucial role in NIH-3T3 Src cells invasiveness by regulating invadosome formation and ECM degradation.

In order to explain the differential role of eIF3 subunits in invadosome formation, we assessed overall protein synthesis in NIH-3T3 Src cells by SunSET. Interestingly we found that eIF3 subunits differentially regulate translation: depletion of eIF3h, eIF3e or eIF3d respectively reduced protein synthesis by 53%, 81% and 75% while the depletion of eIF3k had no effect (Figure 2G). eIF3h is involved in canonical mRNA translation, whereas eIF3d is known to be a driver of non-canonical translation that regulates 20% of mRNAs including mRNAs coding for integrins and metalloproteinases^12,23^. The huge effect of eIF3e depletion on protein synthesis could be explained by its role in both canonical translation through its binding to eIF4g^24^ and non-canonical translation through the regulation of eIF3d expression. These results suggest that both canonical and non-canonical mRNA translation are required for invadosome formation.

### Specific eIF3-dependent mRNAs and product proteins colocalize in invadosomes and control ECM degradation and invasion

Since invadosomes are a translation hot-spot^8^ and β-actin mRNA and protein co-localized with invadosomes (Figure 1A), we hypothesized that specific eIF3-dependent mRNAs could be translated in invadosomes. To identify the mRNAs regulated by eIF3 in invadosomes, we first isolated invadosomes from whole cells by laser capture microdissection associated with next-generation sequencing to establish the invadosome transcriptome (Figure 3A). Thus, we identified 2984 mRNAs enriched in invadosomes compared to whole the cells (Figure 3B). Among the top 30 most significantly enriched mRNAs in invadosomes, a large proportion code for cytoskeletal proteins, such as Acta1/2, Actg2, Tubb2a/b, Tubb3a/b and Tubb5 (Figure 3B and S3A; Table S2), which code for proteins described previously to be involved in invadosomes organization^25–27^.

**Figure 3:**
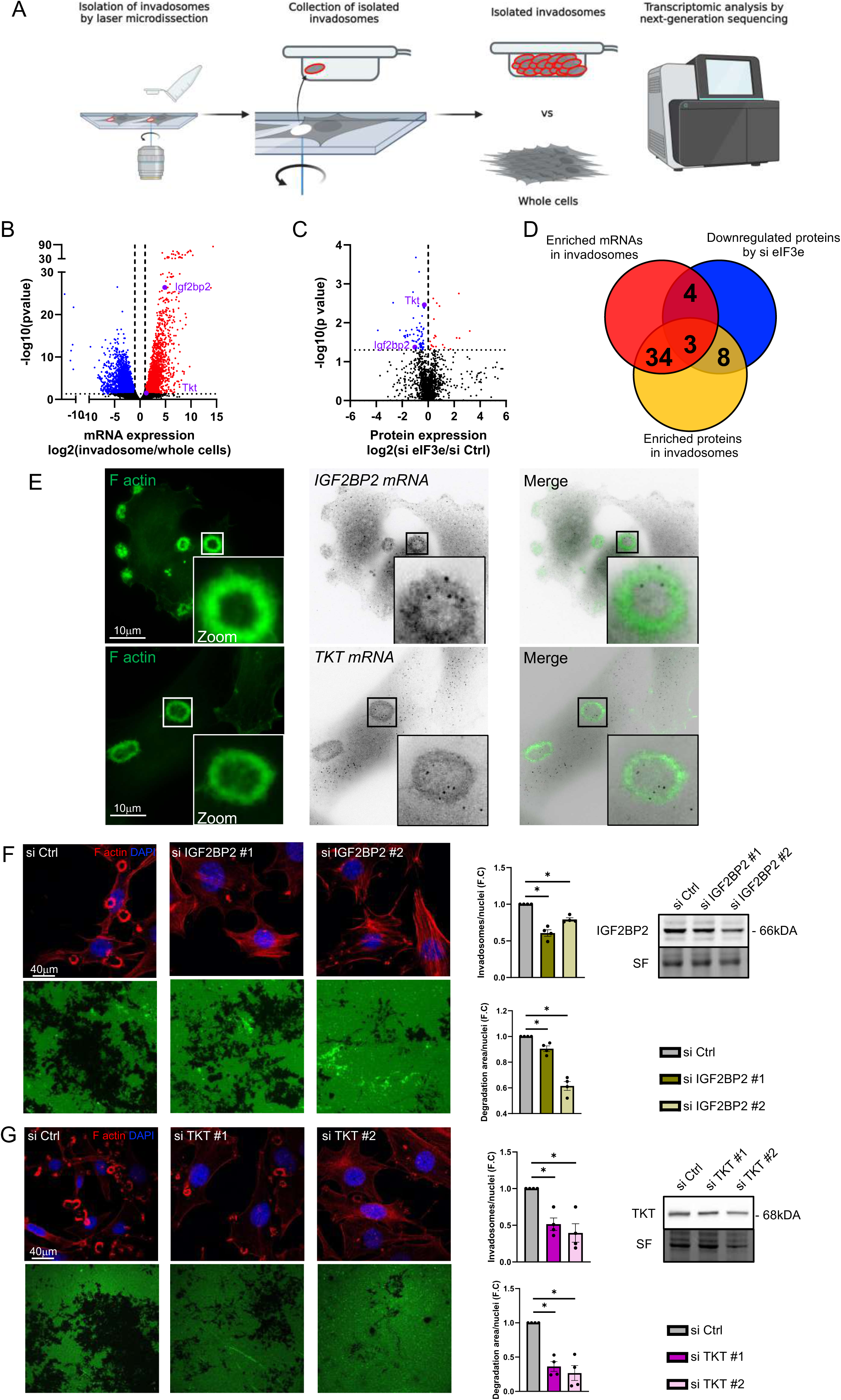
Specific mRNAs and product proteins eIF3-dependent colocalize in invadosomes and control ECM degradation and invasion. (A) Schematic representation of workflow to isolate invadosomes for transcriptomic analysis. (B) Volcano plot displaying mRNAs enriched in invadosomes compared to whole cells, with downregulated mRNAs (blue) and upregulated mRNAs (red). The black ellipse highlights the top 30 of the most significantly enriched mRNAs, selected for gene ontology analysis. IGF2BP2 and TKT expression are highlighted in purple. (C) Volcano plot illustrating proteins downregulated (blue) and upregulated (red) in NIH-3T3 Src (Src) cells treated with siRNA targeting eIF3e compared to control cells. IGF2BP2 and TKT expression are highlighted in purple. (D) Venn diagram demonstrating the overlap of enriched mRNAs and proteins in invadosomes, as well as those downregulated by siRNA targeting eIF3e. (E) Sub-localization of IGF2BP2 and TKT mRNAs via single molecule Fluorescence In Situ Hybridization within invadosomes. Scale bar: 10μm. (F-G) Src cells were transfected with either siRNA control (si Ctrl) or two independent siRNAs targeting IGFBP2 or TKT, then seeded on a fluorescent gelatin matrix. The upper panel presents representative images of invadosomes stained with phalloidin, with the bar graph indicating the number of invadosomes per nucleus. The lower panel displays representative images of the degradation area (depicted in black), with the bar graph quantifying the matrix degradation area per nucleus after 24 hours. Scale bar: 40μm. Error bars (mean ± SEM, *n*=20 fields, four independent experiments, *P<0.05, paired t-test was used for statistical analysis).

By comparative analysis between the proteome^8^ and the transcriptome obtained from isolated invadosomes, we noted a 21% overlap (65 mRNAs and corresponding proteins) between both databases (Figure 3D; Table S3), suggesting that some mRNAs could undergo localized translation in invadosomes. To identify the mRNAs whose translation relies on eIF3, we established the proteome of NIH-3T3 Src cells treated with control or eIF3e siRNA, which represses both canonical and non-canonical translation. Mass spectrometry data analysis revealed the downregulation of 65 proteins upon eIF3e silencing (Figure 3C), including 7 proteins coded by mRNAs we found enriched in invadosomes (Table S4). We then compared the list of mRNAs and corresponding proteins enriched in invadosomes with the list of 65 proteins downregulated after eIF3e silencing; we found 3 common hits (Figure 3D): eIF3e itself, insulin like growth factor 2 mRNA-binding protein 2 (IGF2BP2) and transketolase (TKT). IGF2BP2 is a well-described mRNA-binding protein known to participate in cell motility and formation of cellular protrusions including axon^28^ and invadosomes^29^. TKT is a key enzyme involved in the non-oxidative pentose phosphate pathway and is deregulated in several cancers including liver, pancreatic and colorectal cancers^30^. Recently, it was shown that TKT promotes proliferation, migration and invasion in colorectal cancer cells^31^ but its role in invadosomes is not described in literature. We confirmed that the mRNAs coding for IGF2BP2 and for TKT localize in invadosomes (Figure 3E) and that both proteins are also enriched there (Figure S3B). In order to determine whether IGF2BP2 and TKT are determinant for invadosome formation, we reduced their expression by siRNA in NIH-3T3 Src cells. We demonstrated that depletion of either IGF2BP2 or TKT reduced invadosome formation and matrix degradation activity (Figure 3F and 3G), as well as NIH-3T3 Src cell invasiveness (Figure S3C and S3D), similar to what we found for eIF3e depletion (Figure 2E and S2C).

These results suggest that eIF3-dependent mRNAs, such as *IGF2BP2* and *TKT*, could be translated locally in invadosome to ensure their maintenance for ECM degradation and cellular invasion.

### Src kinase activity enhances canonical and non-canonical eukaryotic translation initiation leading to invadosome formation

To decipher the molecular mechanisms controlled by Src activity and leading to the regulation of eIF3-dependent translation, we performed a comparative Ingenuity Pathway Analysis on the proteome of NIH-3T3 Src cells compared to NIH-3T3 WT cells (Figure 4A, blue dots). This analysis identified that mammalian target of rapamycin (mTOR), ribosomal protein S6 kinase beta-1 (p70S6K) and eIF2 signaling as the most enriched pathways upon Src activation (Figure 4A). Interestingly, mTOR, p70S6K and eIF2 signaling pathways were also enriched in the invadosome proteome we established previously^8^ (Figure 4A, yellow dots). To further identify the kinases whose activity is regulated by Src, we performed a large kinase activity profiling on serine/threonine kinases (STK) and protein tyrosine kinases (PTK) in NIH-3T3 WT and in NIH-3T3 Src cells, treated or not with PP2 (Table S5 and S6). We found that 8 PTKs and 7 STKs (including p70S6K) were modulated by Src activity (Table S5 and S6). A String network analysis showed that 11 out of these 15 Src-activated kinases are linked to mTOR, suggesting that Src is involved in the regulation of mTOR in NIH-3T3 Src cells (Figure S4A). As the phosphoinositide-3-kinase (PI3K)/AKT axis is the most important upstream activator of mTOR signaling^14^ and Src is known to enhance PI3K/AKT activation^32^, we analyzed the phosphorylation status of PI3K/mTOR downstream effectors. We confirmed that the phosphorylation of AKT (S473 and T308), S6K1 (T389) and 4E-BP1 (T37/46) were increased in NIH-3T3 Src cells compared to NIH-3T3 WT cells and were reduced by PP2 or SU6656 treatments (Figure S4B). These results confirm that Src kinase activity regulates upstream and downstream effectors of the mTOR signaling pathway.

**Figure 4:**
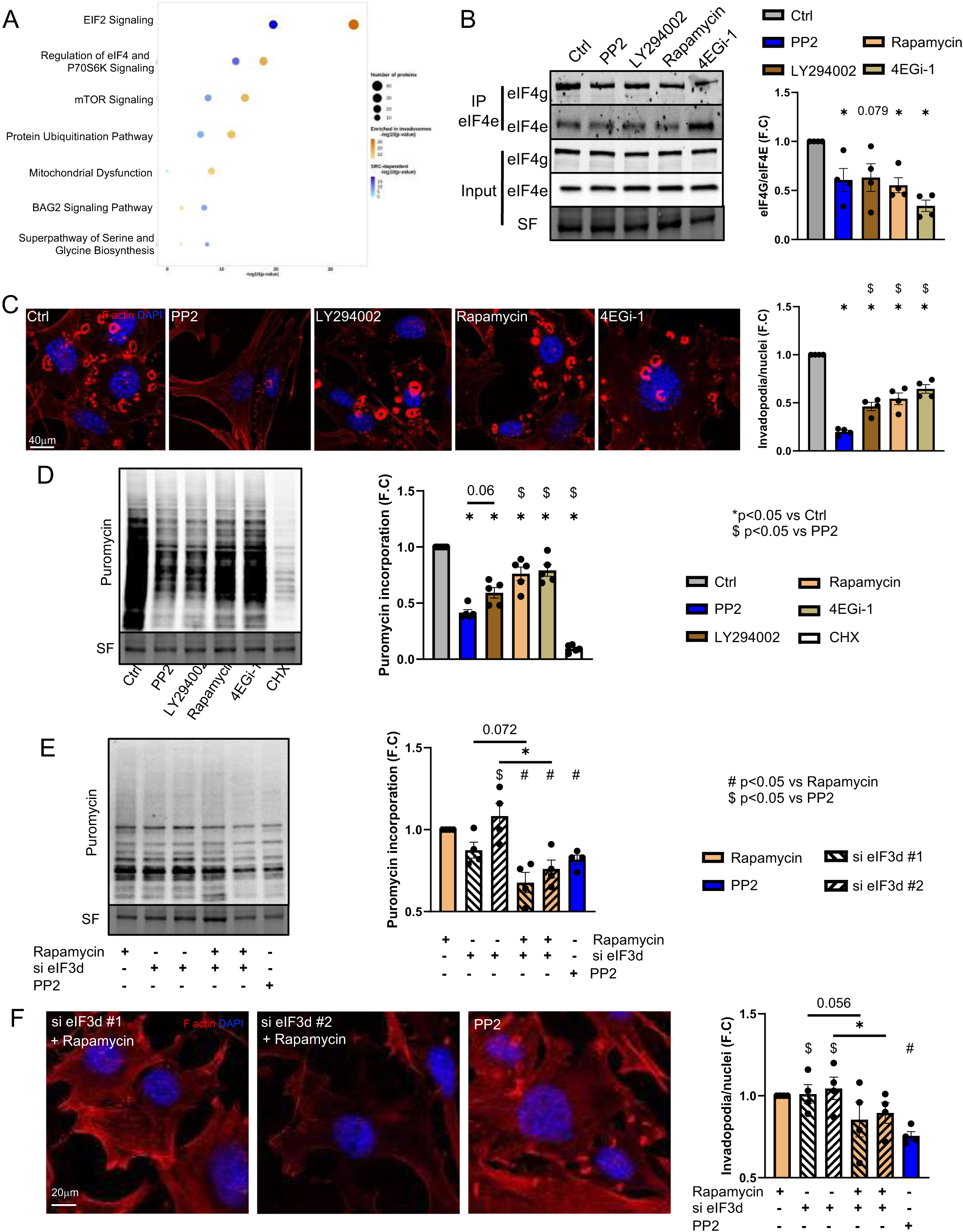
Src kinase activity enhances canonical and non-canonical eukaryotic translation initiation leading to invadosome formation. (A) The bubble plot illustrates the gene set enrichment analysis conducted on the Ingenuity Pathways Analysis database based on the Src-dependent proteome (depicted in blue) and identified enrichment in invadosomes (depicted in yellow). The color and size of the circles indicate the significance level and the numbers of proteins respectively from each enriched pathway. (B-D) NIH-3T3 Src cells underwent treatment with various inhibitors: Src inhibitor (PP2), PI3K inhibitor (LY294002), mTOR inhibitor (Rapamycin), and inhibitor of eIF4e/eIF4g interaction (4EGi-1) during 1 hour. DMSO treatment served as the control group (Ctrl). (B) The left panel presents immunoblot results of immunoprecipitated (IP) and input fractions following eIF4e pull-down. The right panel displays the quantification of the eIF4e/eIF4g ratio in the IP fraction, normalized with respective expression in the input fraction. Error bars (mean ± SEM, four independent experiments, *P<0.05, paired t-test was used for statistical analysis). (C) The left panel exhibits representative images of invadosomes stained with phalloidin, while the bar graph illustrates the number of invadosomes per nucleus. Scale bar: 40μm. Error bars (mean ± SEM, *n*=20 fields, four independent experiments, *P<0.05, paired t-test was used for statistical analysis). (D) SUnSET assay performed on treated NIH-3T3 Src cells. Bar graphs show the quantification of puromycin incorporation. As a negative control, Src cells were incubated with cycloheximide (CHX). (E-F) NIH-3T3 Src cells were transfected with siRNA control (si Ctrl) or with two independent siRNAs targeting eIF3d then treated with rapamycin or PP2 or with DMSO as control during 1 hour. (E) SUnSET assay performed on treated NIH-3T3 Src cells. Bar graphs show the quantification of puromycin incorporation. (F) The left panel exhibits representative images of invadosomes stained with phalloidin, while the bar graph illustrates the number of invadosomes per nucleus. Scale bar: 20μm. Error bars (mean ± SEM, *n*=20 fields, four independent experiments, *P<0.05, paired t-test was used for statistical analysis). Bar graphs for western blot quantification were presented with error bars (mean ± SEM, four to five independent experiments, *P<0.05, paired t-test was used for statistical analysis).

The PI3K/mTOR pathway plays a crucial role in canonical translation initiation by enhancing eIF4e/eIF4g interaction, leading to the association between eIF3 complex to eIF4g, via eIF3e subunit^24^. Thus, we assessed whether Src activation affected eIF4e/eIF4g interaction. We found that Src inhibition by PP2 reduced eIF4e/eIF4g interaction in NIH-3T3 Src cells, assessed by eIF4e immunoprecipitation (Figure 4B). Of note, eIF4g binding to eIF4e was also deduced by treatment with PI3K inhibitor LY294002, mTOR inhibitor rapamycin or 4EGi-1 inhibitor of the eIF4e/eIF4g interaction in NIH-3T3 Src cells, as expected (Figure 4B). We then assessed the effect of these inhibitors on invadosomes. We found that a 1-hour treatment of either inhibitor of canonical translation reduced invadosome formation in NIH-3T3 Src cells, as compare to control (Figure 4C). Interestingly, the inhibition of Src had a stronger effect on invadosomes than any of these treatments (Figure 4C). Similarly, PP2 treatment led to a stronger inhibition of protein synthesis as compared to LY294002, rapamycin and 4EGi-1 (Figure 4D). This suggests that Src could also regulate another translation signaling pathway leading to invadosome formation in NIH-3T3 Src cells. Nevertheless, after a 24-hour treatment, we observed that Src inhibitor and canonical translation initiation inhibitors had a similar inhibitory effect on ECM degradation activity as well as invasion (Figure S4C and S4D).

In order to better understand the differential inhibitory effects on invadosome formation and protein synthesis between Src and PI3K/mTOR inhibitors, we hypothesized that Src could also regulated the non-canonical eIF3d cap-dependent translation pathway, which has been previously demonstrated to regulate determinant proteins (integrins and MMPs)^33^ for invadosome formation and degradation activity. We first showed that the inhibition of Src activity reduced eIF3d expression, supporting the hypothesis that Src could also regulate eIF3d-dependent translation (Figure S4E). We then assess whether the inhibition Src activity has similar effects than the inhibition of both canonical and non-canonical cap-dependent translation. For this, we depleted eIF3d in NIH-3T3 Src cells using siRNAs, and inhibited mTOR with rapamycin. We found that rapamycin treatment and eIF3d depletion had an additional inhibitory effect on protein synthesis and invadosome formation in NIH-3T3 Src cells, which reached a similar inhibitory effect as Src inhibitor (Figure 4E and 4F).

All together these results demonstrate that Src-induced activation of both canonical and non-canonical cap-dependent mRNA translation are determinant for invadosome formation and NIH-3T3 Src cell invasiveness.

### The Src – eIF3h/3e/3d axis is enriched in HCC patients and enhanced the invasiveness of HCC cell line

In order to explore the correlative expression of *Src* and eIF3 subunit genes in human cancers, we used The Human Protein Atlas database (https://www.proteinatlas.org/). We found that high expression of *Src*, *eIF3h*, *eIF3e* and *eIF3d* negatively correlated with patient survival in 4 types of cancer: liver, lung, pancreatic and renal cancers (Figure S5A). Among them, we found a positive correlation between *Src*, *eIF3e* and *eIF3h* gene expression only in liver cancer patients (Figure 5A and S5B), using TCGA data bases (https://portal.gdc.cancer.gov/). Moreover, in patients with hepatocellular carcinoma (HCC), the most common type of liver cancer (http://lifeome.net/database/hccdb/home.html), we found that Src, eIF3h, eIF3e and eIF3d mRNA were increased in the tumor (T) as compared to the non-tumoral (NT) tissue (Figure 5B). We confirmed by immunohistochemistry that eIF3h, eIF3e and eIF3d protein staining is higher in the tumor as compared to the adjacent tissue, while total Src staining was similar (Figure 5C). This suggests that eIF3 have a determinant role in tumor progression in HCC.

**Figure 5:**
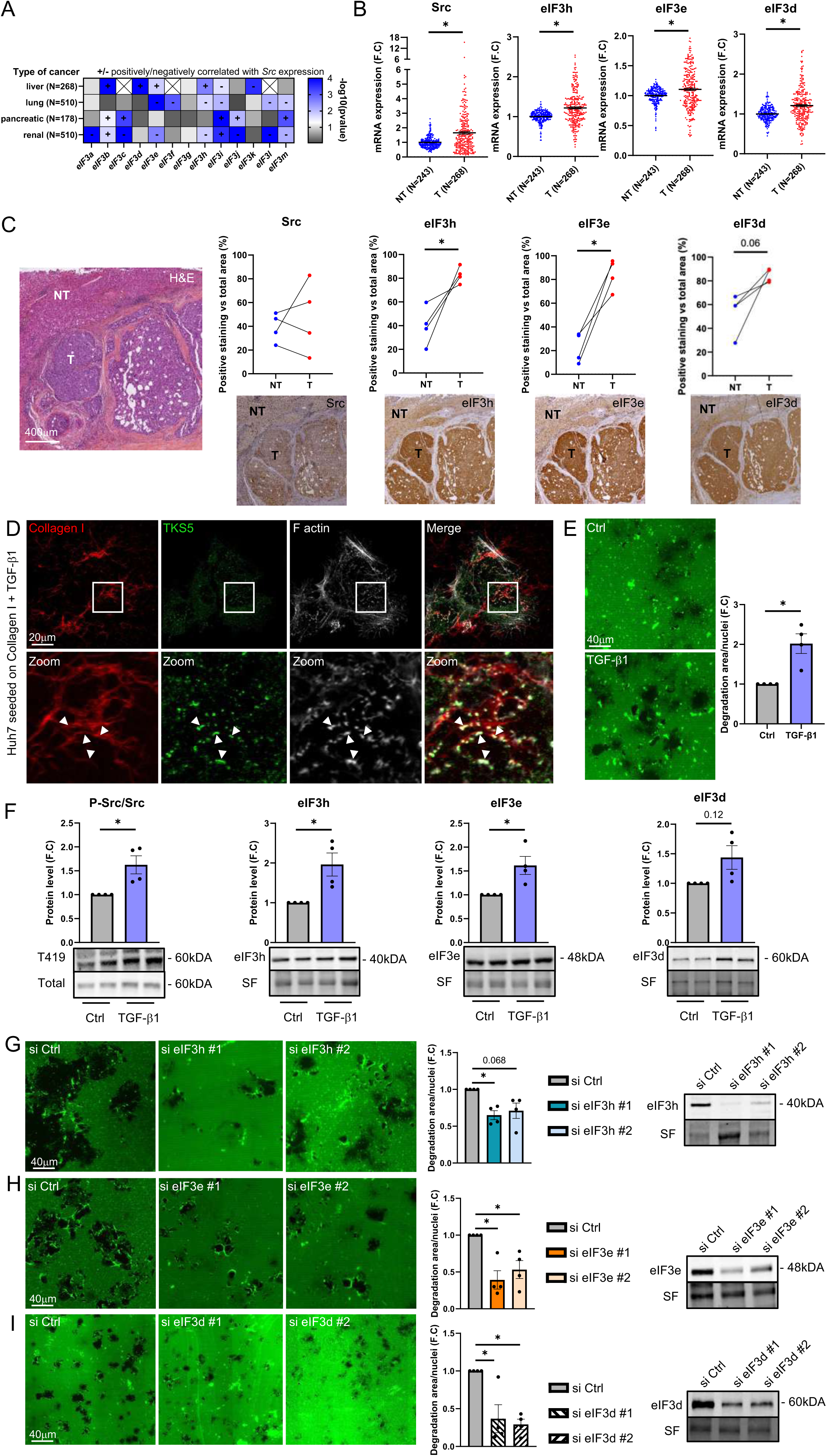
The Src – eIF3h/3e/3d axis is enriched in HCC patients and enhanced the invasiveness of HCC cell line. (A) The heatmap illustrates the correlation between the mRNA expression of Src and eIF3 subunits across four distinct cancer types (liver, lung, pancreatic, and renal). (B) Comparative transcriptomic analysis, from an open database, conducted on tumor (T) versus non-tumor (NT) sections obtained from liver biopsies. (C) Liver biopsies samples from hepatocellular carcinoma (HCC) patients were stained with Hematoxylin/eosin, as well as staining for Src, eIF3h, eIF3e and eIF3d proteins. Bar graphs show positive staining area normalized with total area. Scale bar: 400μm (N=4, *P<0.05, paired t-test was used for statistical analysis). (D) Huh7 cells were treated with Transforming Growth Factor beta 1 (TGF-β1) during 48 hours then seeded on fluorescent collagen type I matrix. TKS5 has been used to reveal linear invadosomes. Heads of arrow indicate the co-localization of TKS5 with actin and collagen I fibers showing linear invadosomes. Scale bar: 20μm. (E) Huh7 cells were cultured on a fluorescent gelatin matrix coated with collagen type I and treated or not with TGF-β1. The left panel presents representative images of the degradation area (depicted in black), with the accompanying bar graph quantifying the matrix degradation area per nucleus after 24 hours. Scale bar: 20μm. Error bars (mean ± SEM, *n*=20 fields, four independent experiments, *P<0.05, paired t-test was used for statistical analysis). (F) Bar graphs illustrate the phosphorylation status of Src and the protein expression levels of eIF3h, eIF3e and eIF3d. Bar graphs for western blot quantification were presented with error bars (mean ± SEM, four to five independent experiments, *P<0.05, paired t-test was used for statistical analysis). (G-I) Huh7 cells were transfected with siRNA control (si Ctrl) or two independent siRNAs targeting eIF3h (G) or eIF3e (H) or eIF3d (I) then cultured on a fluorescent gelatin matrix coated with collagen type I and treated with TGF-β1. The left panel presents representative images of the degradation area (depicted in black), with the accompanying bar graph quantifying the matrix degradation area per nucleus after 24 hours. Scale bar: 40μm. Error bars (mean ± SEM, n=20 fields, four independent experiments, *P<0.05, paired t-test was used for statistical analysis).

In order to examine whether the Src/eIF3 axis could be a main driver for invasiveness in HCC, we used the Huh7 HCC cell line. Huh7 cells are able to acquire an invasive phenotype by the formation of linear invadosomes when they are seeded on type I collagen matrix under the effect of TGF-β1 treatment (Figure 5D)^34^. We confirmed that TGF-β1 treatment stimulated the ECM-degrading activity of Huh7 cells (Figure 5E). Interestingly, TGF-β1 increased Src phosphorylation without affecting total Src levels (Figure 5F). Furthermore, the expression of eIF3h and eIF3e is increased in Huh7 cells treated with TGF-β1 but a slight increase of eIF3d expression was measured (p=0.12) in treated cells (Figure 5F). We then explored the contribution of Src and eIF3 subunits to Huh7 invasiveness. We found that the inhibition of Src activity, or the downregulation of eIF3h or eIF3e or eIF3d with siRNAs prevented the TGF-β1-induced ECM degradation and invasion of Huh7 cells (Figure 5G-5I and S5C-S5G).

All together, these results show that the Src-regulated eIF3h, eIF3e and eIF3d subunits are key players to ECM degradation activity of HCC cells.

## Discussion

Crossing the basement membrane and ECM remodeling are two determining steps enabling invasive cancer cells to migrate and metastasize. In our study, we highlighted a new molecular mechanism downstream of Src kinase and leading to the modulation of protein synthesis by the regulation of the eIF3 complex and both the canonical and non-canonical mRNA translation (Figure 6). We also showed an enrichment in invadosomes of that a variety of eIF3-dependent mRNAs; for some of them, their product protein is also enriched there, such as IGF2BP2 and TKT that play a role in invadosome formation and ECM degradation. We also demonstrated that this new Src/eIF3 signaling pathway is determinant for HCC cell invasiveness.

We and others already demonstrated that local enriched translational machinery has a determinant role for cellular protrusion stabilization^5,8,35^. It has been showed that Src regulates the spatial translation of β-actin through phosphorylation of IGF2BP1 in neuronal outgrowth^6^ which support our hypothesis regarding the potential role of local translation into invadosomes. In accordance with literature, we showed that the regulation of Src activity regulates protein synthesis through canonical pathway following activation of PI3K/mTOR^15^ or ERK/MKK signaling^36^. However, we found that PI3K/mTOR inhibition did not prevent translation and invadosome formation in similar levels as Src inhibition, suggesting that Src signals could control multiple layers of the translational activity. Interestingly, work from Shin *et al*. identified that mTOR/eIF4E axis inhibition promotes cell phenotype switching from proliferative to migratory status through the activation of the non-canonical eIF3d-cap dependent activity^37^. In addition, another study demonstrated that this activity requires the formation of the eIF4g2 (DAP5)/eIF3d dimer promoting cancer cells EMT, invasiveness and metastasis^12^. In our study, we directly connect a potential role of Src activation to the non-canonical eIF3d-cap dependent translation leading to invadosome formation and cell invasion.

As the activation of canonical translation through the PI3K/mTOR pathway is known to promote invadosome formation and metastasis^38^, we demonstrate here that PI3K/mTOR pathway is enriched into invadosomes. We also show that Src-induced PI3K/mTOR activation promotes the interaction between eukaryotic initiation factor 4f subunits eIF4e and eIF4g, necessary for invadosome formation and ECM degradation, which supports the determining role of cap-dependent translation in cell invasiveness. Since eIF4e/eIF4g interaction is required for the cap-dependent translation initiation^14^ and eIF3 complex directly interacts with eIF4g through the eIF3e subunit^24^, our study demonstrated that eIF4e/eIF4g interaction and eIF3e expression have determinant role for Src-induced invadosome formation. Interestingly, the collagen-induced β1 integrin activation increased Src and eIF4g activity in human lung fibroblasts^39^.

This study is the first description of the distinct role of eIF3 subunits in invadosome formation. While depletion of eIF3k has no major effect on invadosomes, depletion of eIF3h, eIF3e and eIF3d strongly impacts both invadosome formation and ECM degradation, associated with lower translation level. In our study, we suggest that eIF3h and eIF3d drive respectively canonical and non-canonical translation while eIF3e, acts at the interconnection of both modes of translation in accordance to its own role in the dynamic of the eIF3 complex, formation^12^, explaining the huge effects of eIF3e depletion in protein synthesis in NIH-3T3 Src cells. A another study confirmed the interest to study eIF3 complex in invadosomes since 11 of the 13 eIF3 subunits were found to interact with p190RhoGAP^40^ a key driver of invadosome formation^41^.

Local translation was first identified in neurons^42^, where mRNA localization and translation at a specific site is involved in various processes such as neurons development, synaptogenesis, experience-dependent plasticity and survival in dendrites and axons. Local translation of mRNAs is also required for cellular protrusion stabilization^35,43^. Many studies were reviewed and deciphered the regulation of mRNA trafficking to their translation nevertheless the exact molecular mechanisms remain unclear^44^. In order to decipher the translatome in invadosomes, we used an innovative method combining transcriptomic and proteomic analysis on isolated invadosomes from the cell body by laser capture microdissection^8^. We were able to highlight 37 mRNAs and their respective coding proteins enriched into invadosomes, suggesting that they could be translated locally. Among these, we identified IGF2BP2 and TKT, whose mRNA translation depends on eIF3, and that are both required for invadosome formation and ECM degradation. IFG2BP2 (or IMP2) was also shown to be involved in dot-like invadosome formation in the triple-negative breast cancer cell line MDA-MB-231^29^. IFG2BP2 is known to enhance mRNA stability and translation, including eIF3 target mRNAs such as c-Myc^45^. Furthermore, a study demonstrated that IFG2BP2 is one of the most enriched proteins in the β-actin mRNA interactome suggesting that IFG2BP2 could stabilize and promote translation of β-actin^46^. Interestingly, TKT, an enzyme involved in non-oxidative pentose phosphate pathway, is highly expressed in several types of cancer and its expression is associated with poor prognosis and increased proliferation and metastasis^30^. Moreover, TKT depletion reduced cell invasiveness in colorectal cancer cells^31^, which is consistent with our finding that TKT is essential for invadosome formation.

Our results showed that Src expression was similar between tumor and non-tumor regions in biopsies from HCC patients. Nevertheless, the activated form of Src (pY416) is positively correlated with tumor grade and associated with poor patients survival in HCC^47^. Furthermore, the activation of pY416-Src remains important an important driver for α_2_-macroglobin-mediated invasion and metastasis in HCC cell lines^48^. We also demonstrated that Src inhibition by PP2 reduced TGF-β1-induced Huh7 invasiveness. More recently, *Yang J et al*. demonstrated that the depletion of Src also reduced both tumor growth and invasion in Huh7 and HepG2 cells^49^. Similarly to Src, the overexpression of several individual eIF3 subunit has been detected in several types of cancer including HCC^50^. We and others demonstrated an increase of *eIF3h* expression in liver cancer including HCC^11,51^. Interestingly, eIF3h overexpression was also found associated with microvascular invasion and advanced tumor-node-metastasis T stage^11^. Our results show that depletion of eIF3h or eIF3e reduced HCC cell line invasiveness in accordance with literature^11^. Overall, our data demonstrate that Src-induced eIF3 expression is a major signaling pathway involved in invasive process, which could represent a potential emerging therapeutic target in HCC.

## Acknowledgements

We thank Prof. Paulette Bioulac-Sage for the expertise in histological analysis. We also thank the Bordeaux Imaging Center (confocal and live cell imaging), Vect’UB (lentivirus production) and Neurocentre Magendie (laser microdissection and transcriptomic facilities) for their support for this study.

## Authors contribution

B.B designed and performed experiments. A.C contributed to experiment realization. S.D.T, J.W.D and A.A.R performed and interpreted mass spectrometry analysis. M.L.B and J.S performed polysome profiling and associated bioinformatical analysis. I.M helped for IHC staining. A.B and C.D contributed respectively to RNAseq and bioinformatic analysis. B.B, A.J, A.B, Y.M and F.S designed the study and F.S lead the study. B.B wrote the manuscript.

---

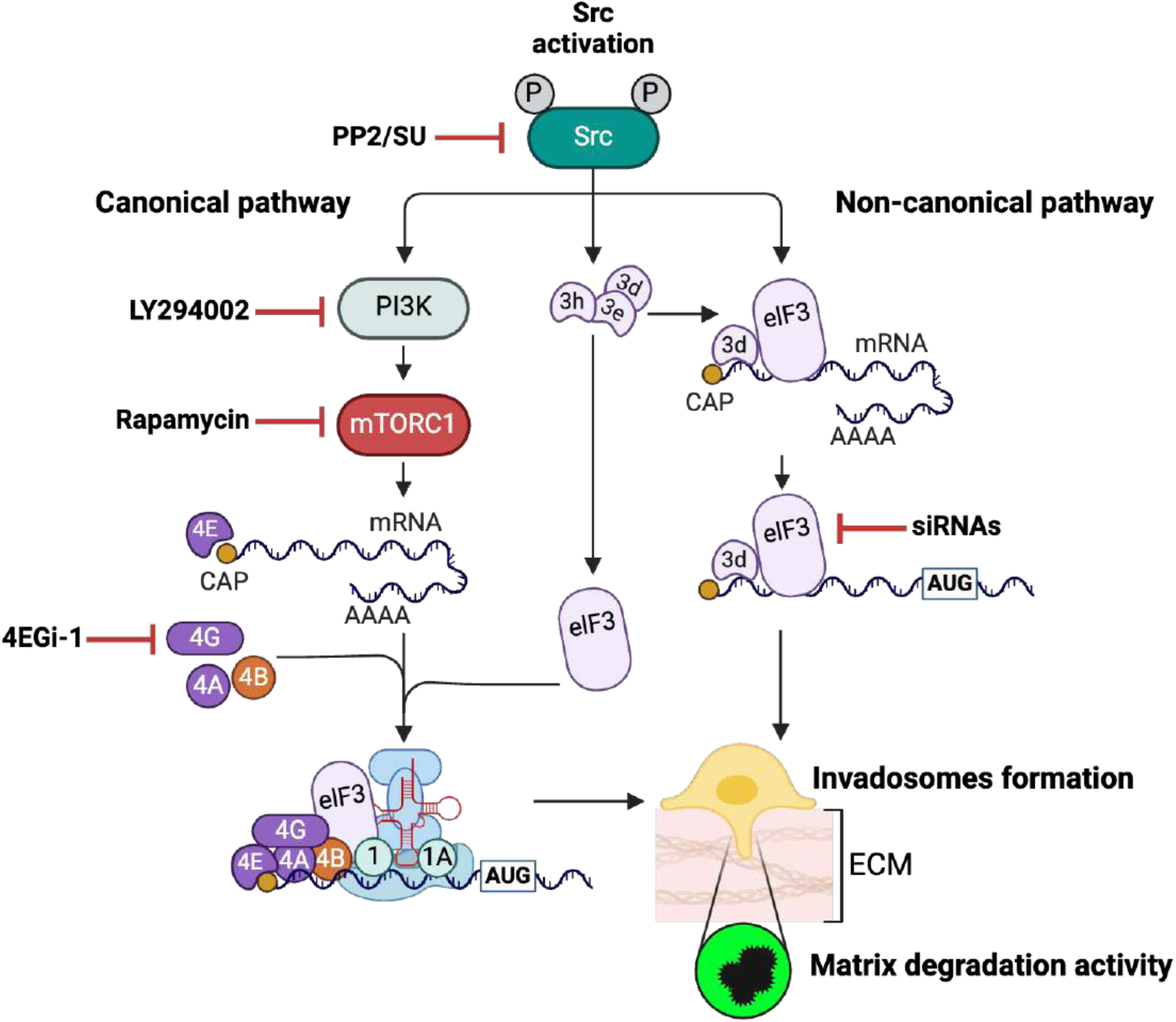

**Figure Supp 1:**
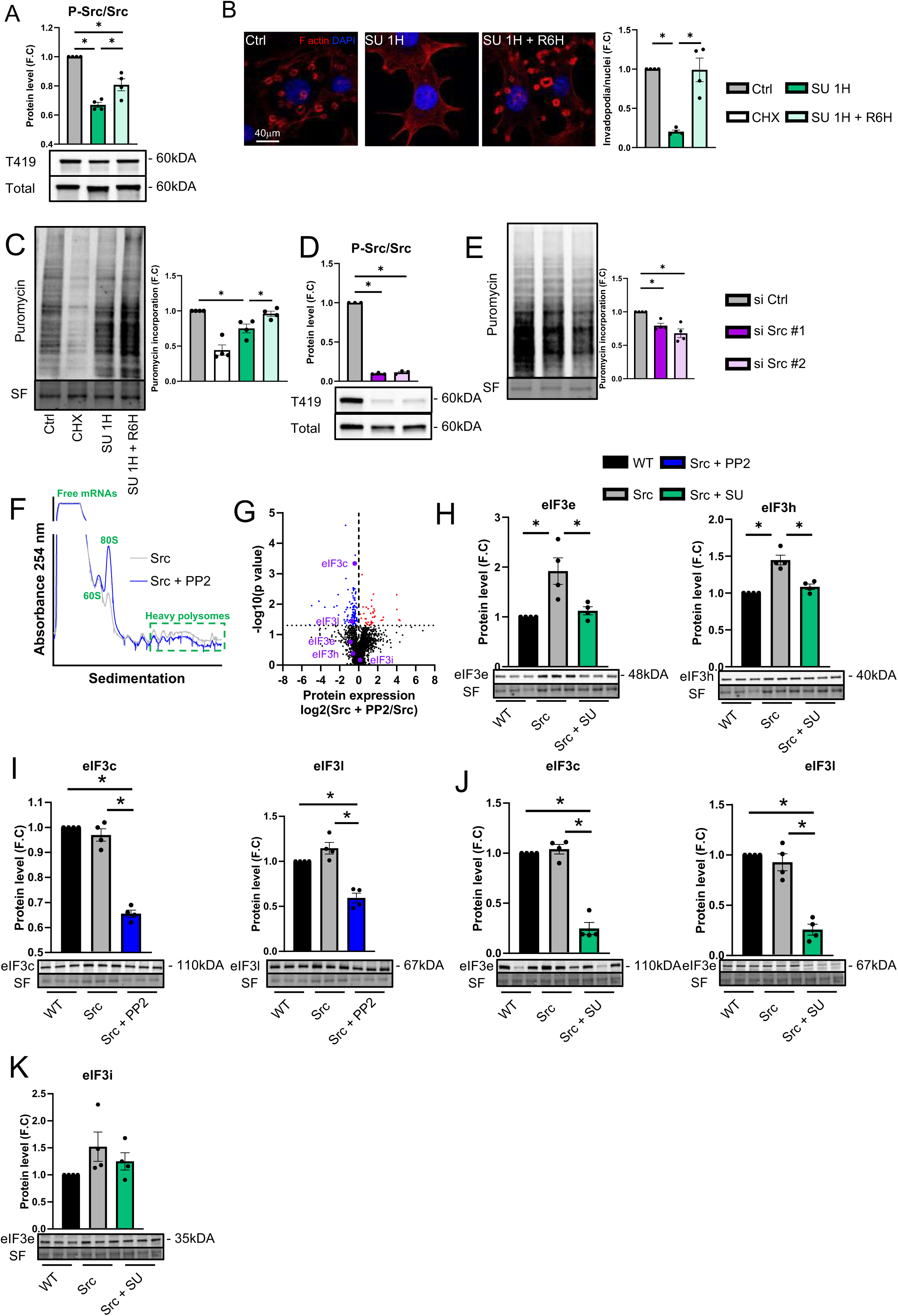
(A-C) NIH-3T3 Src cells were treated with the Src inhibitor SU6656 during 1h (SU 1H) then cells were replaced into fresh medium during recovery period (SU 1H + R6H). DMSO treatment has been used as control group (Ctrl). (A) The bar graph shows the quantification of phosphorylation status of Src. (B) The right panel shows representative images of invadosomes stained with phalloidin; the bar graph shows the number of invadosomes per nuclei. Scale bar: 20μm. Error bars (mean ± SEM, *n*=20 fields, four independent experiments, *P<0.05, paired t-test was used for statistical analysis). (C) SUnSET assay performed on treated Src cells. As a negative control, Src cells were treated with cycloheximide (CHX). Bar graph shows the quantification of puromycin incorporation. (d-e) Src cells were transfected with siRNA control (si Ctrl) or with two independent siRNAs targeting Src. (D) The bar graph shows the quantification of phosphorylation status of Src. (E) SUnSET assay performed on transfected Src cells. Bar graph shows the quantification of puromycin incorporation. (F). Polysome profiles from Src cells treated or not with PP2 during 1h. Absorbance at 254nm is shown as a function of sedimentation. Free mRNAs, the large eukaryotic ribosome subunit (60S), monosome (80S) and heavy polysomes fractions are indicated. (G) Volcano plot of downregulated (blue) and upregulated (red) proteins evaluated by mass spectrometry in Src cells treated or not with Src inhibitor (PP2) during 24 hours. (H-K) Protein expression of eIF3 subunits in WT and Src cells treated or untreated with Src inhibitor (PP2 or SU) for 24 hours. Bar graphs for western blot quantification were presented with error bars (mean ± SEM, four independent experiments, *P<0.05, paired t-test was used for statistical analysis).

**Figure Supp 2:**
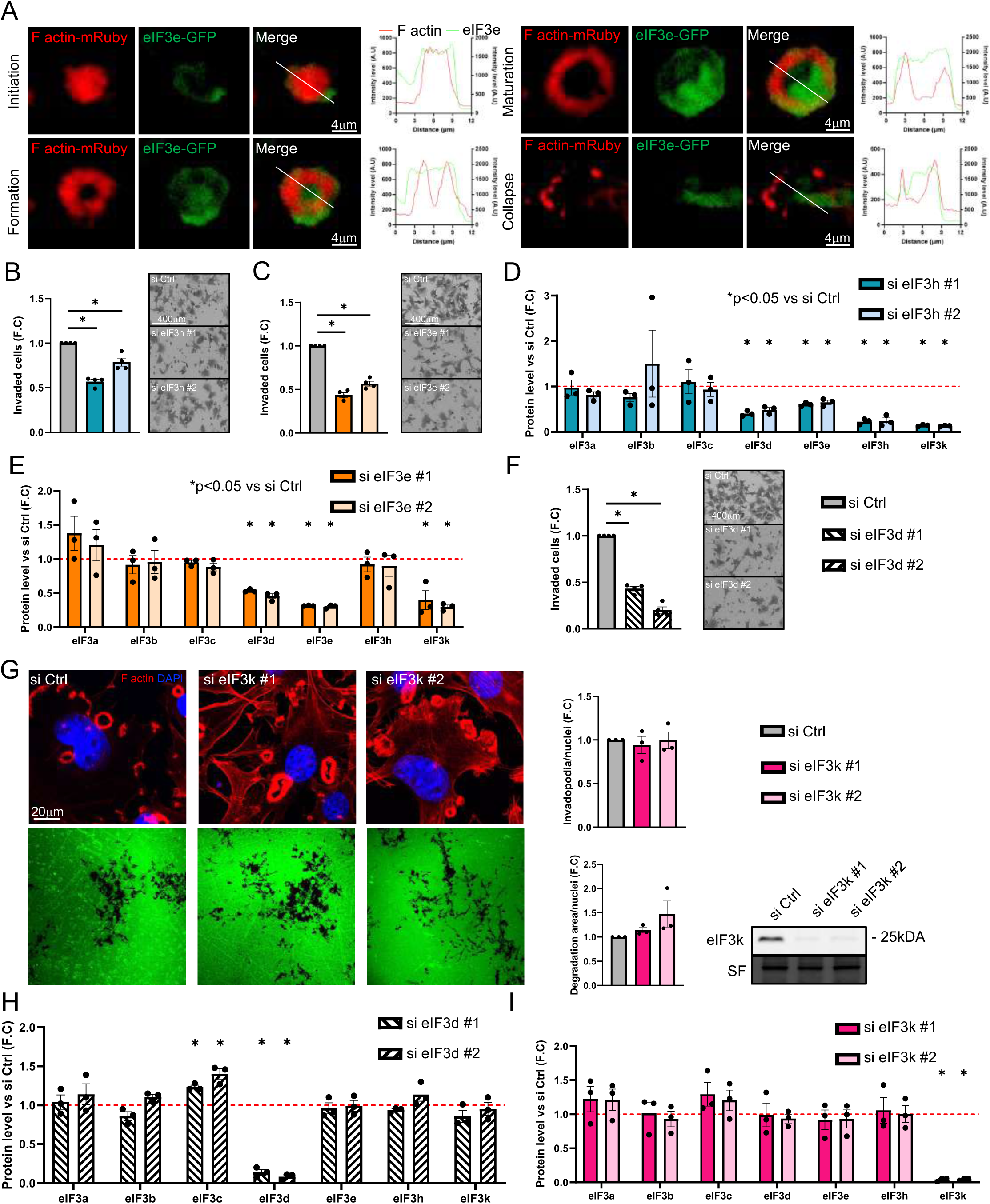
(A) Representative images from time-lapse video microscopy of lifeact-mRuby (red)-expressing NIH-3T3 Src (Src) cells transfected with eIF3e-GFP (green) during invadosome formation (scale bar: 4μm). Bar graphs show F-actin and eIF3e intensity levels as a function of defined distance axis (indicated by white line). (B-I) Src cells were transfected with siRNA control (si Ctrl) or with two independent siRNAs targeting eIF3h or eIF3e or eIF3d or eIF3k. (B-C) Bar graphs show the relative number of invaded NIH-3T3 Src cells treated with siRNAs targeting eIF3h or eIF3e into Boyden chambers. Scale bar: 400μm. Error bars (mean ± SEM, four independent experiments, *P<0.05, paired t-test was used for statistical analysis). (D-E) Protein expression of eIF3 a/b/c/d/e/h/k subunits was evaluated by western blot. Error bars (mean ± SEM, three independent experiments, *P<0.05 compare to si Ctrl, paired t-test was used for statistical analysis). (F) Bar graphs show the relative number of invaded NIH-3T3 Src cells treated with siRNAs targeting eIF3d into Boyden chambers. Scale bar: 400μm. Error bars (mean ± SEM, four independent experiments, *P<0.05, paired t-test was used for statistical analysis). (G) si eIF3k-transfected cells were seeded on a fluorescent gelatin matrix. The upper panel shows representative images of invadosomes stained with phalloidin; the bar graph shows the number of invadosomes per nuclei. The lower panel shows representative images of degradation area (black); the bar graph shows the quantification of matrix degradation area per nuclei after 24 hours. Scale bar: 20μm. Error bars (mean ± SEM, *n*=20 fields, three independent experiments, *P<0.05, paired t-test was used for statistical analysis). (H-I) Protein expression of eIF3 a/b/c/d/e/h/k subunits was evaluated by western blot. Error bars (mean ± SEM, three independent experiments, *P<0.05 compare to si Ctrl, paired t-test was used for statistical analysis).

**Figure Supp 3:**
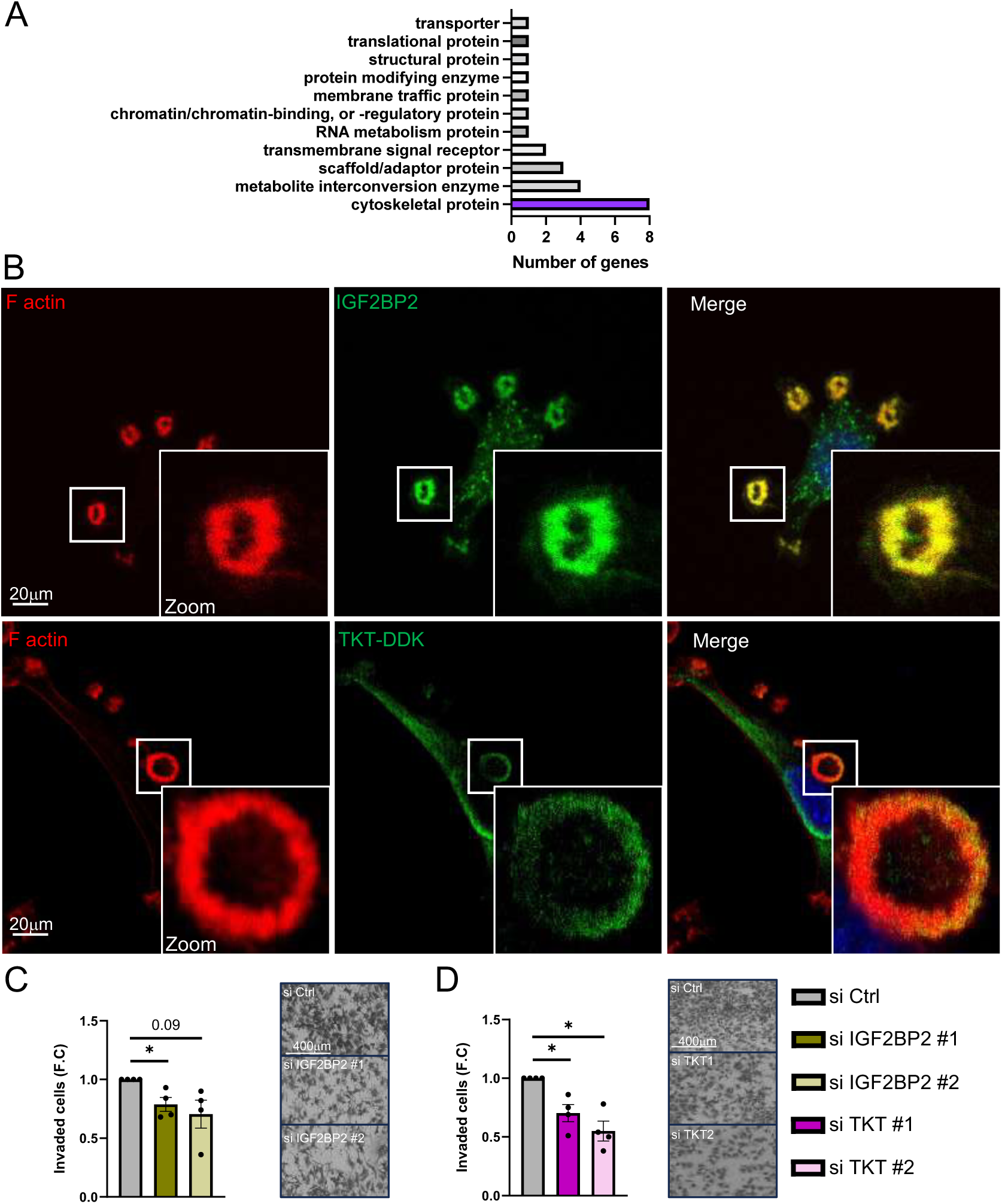
(A) The bar graph shows the protein classes of top 30 mRNAs enriched into invadosomes obtained from gene ontology analysis. (B) Confocal microscopy images of IGFBP2 (green, upper panel), TKT (green, lower panel) and F-actin (red) in NIH-3T3 Src cells. Scale bar: 20μm. (C-D) Bar graphs show the relative number of invaded NIH-3T3 Src cells transfected with siRNA control (si Ctrl) or with two independent siRNAs targeting IGFBP2 (C) and TKT (D) into Boyden chambers. Scale bar: 400μm. Error bars (mean ± SEM, four independent experiments, *P<0.05, paired t-test was used for statistical analysis).

**Figure Supp 4:**
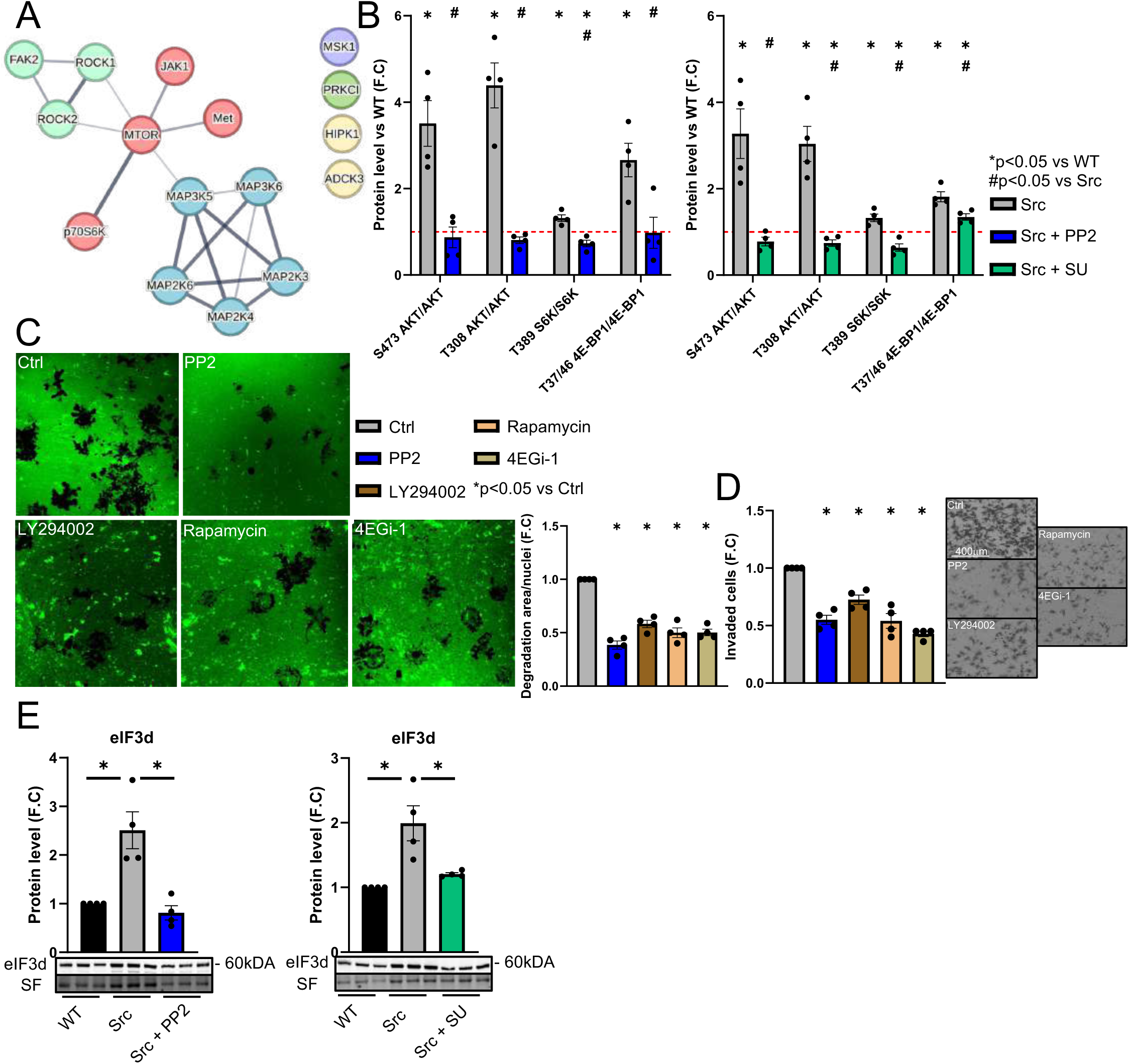
(A) Representation of String network based on kinases regulated by Src activity obtained from Pamgene analysis. (B) NIH-3T3 WT (WT) and NIH-3T3 Src (Src) cells were treated or not with Src inhibitors (PP2 or SU). The phosphorylation status of protein kinase B (AKT), ribosomal protein S6 kinase beta-1 (S6K) and eukaryotic translation initiation factor 4E-binding protein 1 (4E-BP1) were assessed by western blot and normalized to WT cells (dashed red line). Error bars (mean ± SEM, four independent experiments, *P<0.05 compare to WT cells and #P<0.05 compare to Src cells, paired t-test was used for statistical analysis). (C-D) Src cells were seeded then treated with Src inhibitor (PP2), PI3K inhibitor (LY294002), mTOR inhibitor (Rapamycin) and inhibitor of eIF4e/eIF4g interaction (4EGi-1). DMSO treatment has been used as control group (Ctrl). (C) The left panel exhibits representative images of degradation area, while the bar graph illustrates the number of invadosomes per nucleus. Scale bar: 20μm. Error bars (mean ± SEM, *n*=20 fields, four independent experiments, *P<0.05, paired t-test was used for statistical analysis). (D) Bar graph shows the relative number of invaded Src cells treated with inhibitors into Boyden chambers. Scale bar: 400μm. Error bars (mean ± SEM, four to five independent experiments, *P<0.05 compare to Ctrl, paired t-test was used for statistical analysis). (E) Protein expression of eIF3d in Src cells treated with Src inhibitors (PP2 or SU). Error bars (mean ± SEM, four independent experiments, *P<0.05, paired t-test was used for statistical analysis).

**Figure Supp 5:**
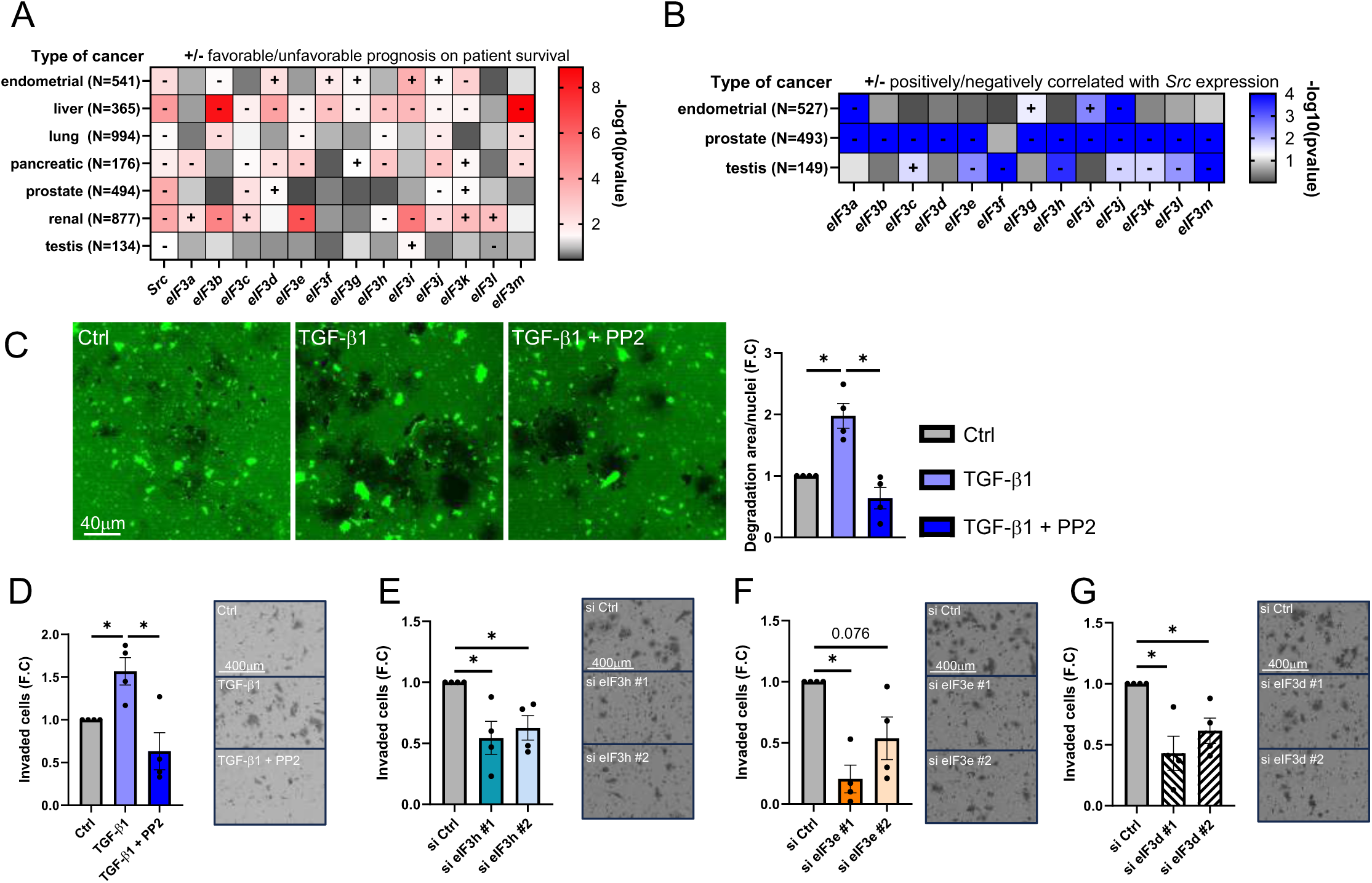
(A) Heat map shows the correlation between survival prognosis status with high expression level of targeted mRNAs in selected types of cancers (B) Heat map shows the mRNA expression correlation between Src and eIF3 subunits in three selected types of cancers (endometrial, prostate and testis). (C) Huh7 cells were seeded on a fluorescent gelatin matrix coated with collagen type I and treated or not with TGF-β1 alone or in combination with Src inhibitor (TGF-β1 + PP2). The left panel depicts representative images of the degradation area (depicted in black), with the accompanying bar graph quantifying the matrix degradation area per nucleus after 24 hours. Scale bar: 20μm. Error bars (mean ± SEM, n=20 fields, four independent experiments, *P<0.05, paired t-test was used for statistical analysis). (D) Huh7 cells were seeded on Boyden chambers coated with collagen type I and treated or not with TGF-β1 associated or not with Src inhibitor. Bar graph shows the relative number of invaded cells. Scale bar: 400μm. Error bars (mean ± SEM, four independent experiments, *P<0.05 compare to Ctrl, paired t-test was used for statistical analysis). (E-G) Huh7 cells were transfected with siRNA control (si Ctrl) or with two independent siRNAs targeting eIF3h or eIF3e or eIF3d treated with TGF-β1 then seeded on Boyden chambers coated with collagen type I. Bar graph shows the relative number of invaded cells. Scale bar: 400μm. Error bars (mean ± SEM, four independent experiments, *P<0.05 compare to si Ctrl, paired t-test was used for statistical analysis).

