## Supplementary material for "Src promotes tumor cell invasion by hijacking the translation machinery": Material and methods: Material and methods.docx

**Cell culture**

NIH-3T3 WT, NIH-3T3-Src cells were a generous gift from Sara A. Courtneidge (Burnham Institute for Medical Research, LaJolla, CA). NIH-3T3 WT, NIH-3T3-Src and Huh7 cells were maintained in Dulbecco’s modified Eagle’s medium (DMEM) with 4.5 g/L glucose Glutamax-I (Invitrogen) supplemented with 10% fetal calf serum (SigmaAldrich). All the cell lines were confirmed for the absence of mycoplasma by PCR.

**Transfection**

The plasmids eIF3h-GFP (Vector Builder #230803) and eIF3e-GFP (Vector Builder #221031) were transduced in NIH-3T3-Src and NIH-3T3-Src Lifeact-mRuby cells by lentiviral infection. Transduced cells were sorted (BD FACSARIA III cell sorter) and re-plated in culture to be used. TKT-DDK plasmid was purchased from Origen and transfected (1μg) using JetPrime (PolyPlus transfection) following the provider’s instructions. siRNA oligonucleotides (50nM) were transfected by Lipofectomine RNAiMax (Invitrogen). siRNAs were listed in Table S7.

**Western Blot**

Cells were washed twice in ice-cold PBS and scraped into ice-cold RIPA lysis buffer (Sigma) completed with Complete and PhoStop inhibitor (Roche Applied Science) and lysed for 30 min at 4 °C. Lysates were then clarified by centrifugation at 13,000 g for 10 min at 4 °C. Protein concentration was measured using Pierce BCA protein assay (ThermoFisher). Proteins (20μg) were boiled and separated using TGX Stain-Free FastCast gels Acrylamide 10% (Biorad) at 120V for 90min. Proteins were transferred into nitrocellulose membrane (Biorad) using Trans-blot Turbo System (Biorad). Membranes were blocked in 5% BSA TBS-T (50 mM Tris, 150 mM NaCl and 0.1% Tween-20 supplemented with 5% BSA) during 1h at room temperature before to be incubated with primary antibodies (1/2000 in 5% BSA TBS-T) during overnight at 4°C. After 3 washes in TBS-T, membranes were incubated with fluorochrome-conjugated secondary antibodies (1/5000 in 5% BSA TBS-T) during 1h at room temperature. After 3 washes in TBS-T, membranes were exposed to right excitation sources using ChemiDoc MP Imaging System (Biorad).

**Antibodies and reagents**

List of antibodies for western blotting, immunofluorescence, immunohistochemistry and immunoprecipitation is provided in Table S8. For pharmacological approach, cells were treated with PP2 (5μM, Sigma), SU6656 (5μM, Sigma), LY294002 (10μM, Cell Signaling), rapamycin (100nM, Sigma) and 4EGi-1 (50μM, Sigma). Huh7 cells were stimulated with 3ng/ml of TGF-β1 (Sigma-Aldrich).

**eIF4E immunoprecipitation and eIF4G blot analysis**

1mg of cell lysate were incubated with 4 μg of eIF4E antibody during overnight at 4°C with constant agitation. The same amount of mouse IgG has been used as control. Protein G Dynabeads (Life Technologies) were washed twice with PBS and once with RIPA lysis buffer before incubation for 3h at 4 °C with samples containing antibodies or IgG. After 5 time washing with RIPA buffer, precipitated proteins were eluted in 2× LDS sample buffer containing 50 mM dithiothreitol (DTT) and boiled at 95 °C for 10 min before analysis via SDS–PAGE electrophoresis and western blotting.

**Sunset assay**

Cells were treated with puromycin (10mg/ml) during 10min at 37°C then washed twice in ice-cold PBS for protein extraction as described above in Western Blot section. For negative control we pre-treated cells with the translation inhibitor cycloheximide (35μM) during 10min at 37°C.

**Degradation assay**

Sterile coverslips were coated with Oregon Green 488-conjugated Gelatin (Invitrogen) for 20 min, then fixed with 0.5% glutaraldehyde (Sigma) for 40 min. After twice washing with PBS, coated-coverslip were placed into 24 wells/plate and cells were seeded (20K/well). Degradation area was measured 24h after seeding and a total of 20 images were acquired for each condition and normalized by number of nuclei in each image. Quantification has been done using ImageJ software.

For HUH7 degradation assays, cells were pre-treated or not with Transforming Growth Factor beta 1 (TGF-β1, 3ng/ml) during 48h as previously described^1^. The pre-coated gelatin coverslips were incubated with type 1 collagen (0.5mg/ml) diluted in DPBS during 4 hours at 37°C. Excess of collagen was removed then cells were seeded (40K/well) treated or not with TGF-β1 (3ng/ml) during 24h.

**Invasion assay**

Boyden chambers (8.0µm pore size, Falcon) were coated with 150µl of Matrigel (80µg/ml) in a 24-well plate. For HUH7 cells, cells were treated and Boyden chambers were also coated with type 1 collagen as described before in degradation assay section. Cells were pre-treated with mitomycin (10μg/mL) during 2h at 37°C before to be seeded in fresh serum-free medium (50K/insert). As a chemoattractant, 20% FBS complete medium was used in the lower part of the transwell in the 24-well plate. After 24 h, the media was carefully removed, cells were fixed with 4% PFA (Electron Microscopy Sciences) for 10 min and stained with crystal violet (Sigma-Aldrich) for 10 minutes at room temperature. After twice washing with distilled water, cells in the upper part of the transwell membrane were removed by wiping with a cotton swab. The crystal violet has been eluted by adding 400μL of 33% acetic acid into each insert and shaking for 10min. The absorbance of eluted crystal violet has been measured at 590nm and the number of invaded cells has been quantified following established standard curve.

**Polysome profiling + Bioinformatical analyses**

NIH-3T3-Src cells were cultivated in 15 mm culture dishes and treated with PP2 at sub-confluence as described above. Cells were treated with 100 µM cycloheximide (CHX) for 5 minutes before being harvested on ice. Cells were washed twice with cold PBS with 100 µM cycloheximide and lysed in 350 μL hypotonic lysis buffer (5 mM Tris-HCl pH 7.5, 2.5 mM MgCl2, 1.5 mM KCl, 100 μg/ml cycloheximide, 2 mM DTT, 0.5% Triton X-100, 0.5% sodium deoxycholate and 1mM Ribonucleoside vanadyl complexes). After normalizing the RNA concentration, lysates were loaded onto a 5 mL linear 10%-45% gradient, and centrifuged at 45,000 rpm for 45 minutes at 4°C. The absorbance at 254 nm was continuously recorded using an ISCO fractionator. For mRNA translation analysis (adapted from Liang et al. NAR 2018 PMID: 29069469), 250 μL of lysates were loaded onto a 5 mL discontinuous 5%-34%-55% sucrose gradient (20 mM HEPES-KOH pH 7.6, 100 mM KCl, 5 mM MgCl2) and centrifuged at 45,000 rpm (SW 55 Ti rotor, Beckman Coulter, Inc.). Fractions containing mRNAs associated to more than 3 ribosomes, were pooled and purified using TRIzol-LS (Thermo Fisher Scientific) in parallel to 50 μL of the total lysate. RNA were treated with DNAse-I using RNA Clean & Concentrator kit (Zymo Research). The quality (DV200> 60%) and the quantity (concentration) of all mRNA samples were evaluated using a Fragment Analyser (Agilent) prior sequencing.

Changes in translational efficiency were quantified using analysis of partial variance as implemented in the anota2seq R/Bioconductor package version 1.24.0 (Oertlin et al, 2019). Both preprocessing and differential expressions of cytosolic and polysome‐associated RNA was assessed using the anota2seq package after a correlation and PCA initial exploration. Sequencing batch was considered for correction while TMM-log2 normalization was utilized in the built-in parameters. For such analyses, Benjamini–Hochberg correction was used to account for multiple testing with a 95% FDR threshold. A random variance model was used to increase statistical power (Wright & Simon, 2003; Larsson et al, 2010). All regulatory modes were considered for the analysis.

**Immunofluorescence and imaging**

Cells were fixed using 4% PFA for 10min at room temperature then rinsed 3 times with PBS. Cells were permeabilized with 0.1% Triton X-100 (Sigma) for 10 min at room temperature before being rinsed twice with PBS. Cells were then incubated with primary antibodies (1:100 diluted in 4% BSA-PBS) for 40 min at room temperature, rinsed 3 times with PBS and incubated with secondary antibodies (1:100 diluted in 4% BSA-PBS) for 30 min at room temperature. Nuclei were stained with DAPI (1:200 diluted) and actin was stained with phalloidin (1:200 diluted). Coverslips were mounted on microscope slides using Fluoromount-G mounting media (SouthernBiotech) and were imaged using SP5 confocal microscope (Leica). Images were analyzed using ImageJ software. Invadosomes counting was performed on F-actin-stained cells 24h after seeding and a total of 20 images were acquired for each condition and normalized by number of nuclei in each image. Quantification has been done using ImageJ software.

**Co-staining β-actin mRNA and coding protein**

Coding sequences for tandem-dimer stdMCP-stdGFP (Add Gene #98916) and HaloTag-β-actin (Add Gene #102718) were cloned into the lentivirus expression vector then used to transfect NIH-3T3 Src cells as previously described^2^. To visualize β-actin protein, we incubated cells with Halo-ligand JF549 (Promega) at final concentration of 1μM during 30 min following provider’s instructions before cell fixation and imaging as described see above.

**Single Molecule Fluorescence In Situ Hybridization**

Single Molecule Fluorescence In Situ Hybridization (smFISH) was performed as described (Tsanov et al., 2016). Briefly, NIH-3T3-Src cells expressing LifeAct-GFP were fixed in 4% paraformaldehyde and permeabilized in 70% ethanol. Cells were then hybridized using two types of probes: (i) 24 unlabeled primary probes containing both an Igf2bp2 or a Tkt mRNA targeting sequence and a shared sequence (FLAP); (ii) a secondary probe conjugated to two Cy3 moieties, pre-hybridized in vitro to the primary probes via the FLAP sequence. Probes for Igf2bp2 and Tkt mRNAs were designed using Oligostan R-Scrip (<https://bitbucket.org/muellerflorian/fish_quant/src/master/Oligostan/>). All probes were purchased from Integrated DNA Technologies (Belgium); the sequences are listed in Table S9. Nuclei were stained with Hoechst and cells were mounted in CitiFluor (Electron Microscopy Sciences, USA). Image acquisition was performed on a Zeiss Axio Imager Z2 equipped with an Xcite 120LED Boost excitation source and a Orca Flash 4.0 V2+ Hamamatsu camera, using a 63X Plan Apochromat 1.4 NA oil objective controlled by Zen Software. Images were acquired as 0.24 µm z-stacks and processed with ImageJ.

**Video microscopy**

Cells were seeded in a bottom-glass dish (µ-Dish 35mm, Ibidi). Cells were imaged every 2 min for 4h using W1 LiveSR spinning disk microscope at the Bordeaux Imaging Center. SPI-488 and SPI-561 were used to image eIF3H-GFP or eIF3E-GFP and Lifeact-mRuby respectively.

**Laser microdissection**

Invadosomes were microdissected from PFA fixed NIH-3T3 Src Lifeact-mRuby cells with a PALM type 4 (Zeiss) automated laser micro dissector.

**Label-free quantitative proteomics**

Three independent biological replicates on total protein extracts were compared by label-free protein quantification. Proteins were loaded on a 10% acrylamide SDS-PAGE gel and proteins were visualized by Colloidal Blue staining. Migration was stopped when samples had just entered the resolving gel and the unresolved region of the gel was cut into only one segment. Each SDS-PAGE band was cut into 1 mm x 1 mm gel pieces. Gel pieces were destained in 25 mM ammonium bicarbonate (NH_4_HCO_3_), 50% Acetonitrile (ACN) and shrunk in ACN for 10 min. After ACN removal, gel pieces were dried at room temperature. Proteins were first reduced in 10 mM dithiothreitol, 100 mM NH_4_HCO_3_ for 60 min at 56°C then alkylated in 100 mM iodoacetamide, 100 mM NH_4_HCO_3_ for 60 min at room temperature and shrunk in ACN for 10 min. After ACN removal, gel pieces were rehydrated with 50 mM NH_4_HCO_3_ for 10 min at room temperature. Before protein digestion, gel pieces were shrunk in ACN for 10 min and dried at room temperature. Proteins were digested by incubating each gel slice with 10 ng/µl of trypsin (V5111, Promega) in 40 mM NH_4_HCO_3_, rehydrated at 4°C for 10 min, and finally incubated overnight at 37°C. The resulting peptides were extracted from the gel by three steps: a first incubation in 40 mM NH_4_HCO_3_ for 15 min at room temperature and two incubations in 47.5% ACN, 5% formic acid for 15 min at room temperature. The three collected extractions were pooled with the initial digestion supernatant, dried in a SpeedVac, and resuspended with 0.1% formic acid for a final concentration of 0.05 µg/µL. NanoLC-MS/MS analysis were performed using an Ultimate 3000 RSLC Nano-UPHLC system (Thermo Scientific, USA) coupled to a nanospray Orbitrap Fusion™ Lumos™ Tribrid™ Mass Spectrometer (Thermo Fisher Scientific, California, USA). Each peptide extracts were loaded on a 300 µm ID x 5 mm PepMap C_18_ precolumn (Thermo Scientific, USA) at a flow rate of 10 µL/min. After a 3 min desalting step, peptides were separated on a 50 cm EasySpray column (75 µm ID, 2 µm C_18_ beads, 100 Å pore size, ES903, Thermo Fisher Scientific) with a 4-40% linear gradient of solvent B (0.1% formic acid in 80% ACN) in 57 min. The separation flow rate was set at 300 nL/min. The mass spectrometer operated in positive ion mode at a 2.0 kV needle voltage. Data was acquired using Xcalibur 4.4 software in a data-dependent mode. MS scans (m/z 375-1500) were recorded at a resolution of R = 120000 (@ m/z 200), a standard AGC target and an injection time in automatic mode, followed by a top speed duty cycle of up to 3 seconds for MS/MS acquisition. Precursor ions (2 to 7 charge states) were isolated in the quadrupole with a mass window of 1.6 Th and fragmented with HCD@28% normalized collision energy. MS/MS data was acquired in the ion trap with rapid scan mode, a 20% normalized AGC target and a maximum injection time in dynamic mode. Selected precursors were excluded for 60 seconds. Protein identification was done in Proteome Discoverer 2.5. Mascot 2.5 algorithm was used for protein identification in batch mode by searching against a UniProt *Mus musculus* protein database (55,334 entries, release December 7, 2022 ; https://www.uniprot.org/ website). Two missed enzyme cleavages were allowed for the trypsin. Mass tolerances in MS and MS/MS were set to 10 ppm and 0.6 Da. Oxidation (M) and acetylation (K) were searched as dynamic modifications and carbamidomethylation (C) as static modification. Raw LC-MS/MS data were imported in Proline Studio for feature detection, alignment, and quantification^3^. Proteins identification was accepted only with at least 2 specific peptides with a pretty rank=1 and with a protein FDR value less than 1.0% calculated using the “decoy” option in Mascot. Label-free quantification of MS1 level by extracted ion chromatograms (XIC) was carried out with parameters indicated previously^4^. The normalization was carried out on median of ratios. The inference of missing values was applied with 5% of the background noise. The mass spectrometry proteomics data have been deposited to the ProteomeXchange Consortium via the PRIDE^5^ partner repository with the dataset identifier PXD000000 and PXD000000.

**RNA extraction**

Isolation of the total RNA from each sample was performed with Direct-zol RNA miniprep (Zymo) following the supplier procedures. The concentration of RNA was measured by spectrophotometry (NanoDrop 2000, ThermoFisher) for each sample. RNA quality was checked using Lab Chip GX touch HT (PerKin Elmer) and RNA chip DNA 5K/RNA CZE (Perkin Elmer).

**Libraries synthesized**

cDNA libraries were synthesized using 500 ng of RNA from each sample with the Illumina Stranded mRNA Prep Ligation (Illumina) kit following the supplier procedures. Briefly, the first step consists on capturing mRNAs with magnetic beads targeting their poly-A tail. Then, mRNAs were fragmented, reverse transcribed into double strand cDNA and 3’ adenylated adapters and anchors were ligated to both ends. Finally, the cDNA library was amplified by PCR with a number of PCR cycles adapted to the start material quantity. Libraries quality was checked with Xmark chip on the Lab Chip GX touch HT (Perkin Elmer). After quantification of individual libraries by q-PCR (Roche LC480) using the NEBNext library quant kit (New England BioLabs), all the samples were pooled in an equimolar manner. The quality and quantity of the pool were checked by qPCR (Roche LC480) with the same kit and Lab Chip GX touch HT (Perkin Elmer) before sequencing. mRNA sequencing was performed using Nextseq 2000 Illumina (paired-end 2x100 bp) with a minimum of 30 million reads per sample in the PGTB facility (INRAE - Pierroton).

**Human database**

The Human database related to patient survival and expression level of targeted genes were extracted from TCGA Pan-Cancer Atlas 2018.

**Patient samples**

Liver FFPE samples were clinically and histologically reviewed by Prof. Bioulac-Sage. Liver FFPE slices were performed using microtome instrument then immunochemistry was performed with antibodies listed in Table S8.

**Bioinformatics analysis**

Proteins and mRNAs lists provided in Table S1-7 were used for Ingenuity Pathways Analysis (Qiagen), partner interaction network (STRING)^6^ and Gene Ontology classification from the PANTHER database ([https://pantherdb.org/).](https://pantherdb.org/)

**Statistical analysis**

Statistical significance was determined using Student’s t-test and calculated using GraphPad Prism software.
