## Supplementary material for "Src promotes tumor cell invasion by hijacking the translation machinery": Tables: Tables.docx

| **Translational proteins** | | | |
| --- | --- | --- | --- |
| **Src/WT** | **Src+PP2/Src** | **Enriched in invadosomes** | |
| Cars1; Eef1a1; Eif2b3; **Eif3e**; **Eif3f**; **Eif3g**; **Eif3h**; Psmd7; Sars1 | Aars1; Cars1; Eef2; Eif3a; Eif3b; **Eif3c**; **Eif3l**; Eif4b; Eif4g1; Eif5b; Gars1; Gspt1; Iars1; Kars1; Lars1; Rpl19; Tars1; Vars1 | Aimp1; Cars1; Dars; Eef1a1; Eef1a2; Eef1d; Eef2; Eftud2; Eif2a; Eif2s1; **Eif3c**; **Eif3e**; **Eif3f**; **Eif3g**; **Eif3h**; **Eif3i**; Eif3k; **Eif3l**; Eif4b; Eif5a; Elavl1; Eprs; Farsa; Fxr1; Rars; Rpl10; Rpl10a | Rpl12; Rpl22; Rpl23; Rpl26; Rpl28; Rpl30; Rpl31; Rpl34; Rpl35a; Rpl36a; Rps11; Rps12; Rps13; Rps14; Rps15a; Rps16; Rps19; Rps20; Rps26; Rps27; Rps3; Rps3a; Rps4x; Rpsa; Vars; Yars |

Table S1: Translational proteins classification by Gene Ontology analysis of proteins regulated by Src and enriched in invadosomes.

**Bold** = commonly regulation of translational factors

| **ENSEMBL** | **Gene name** | **Protein name** | **Protein class** |
| --- | --- | --- | --- |
| **ENSMUSG00000031972** | **Acta1** | **Actin, alpha skeletal muscle** | **cytoskeletal protein (PC00085)** |
| **ENSMUSG00000035783** | **Acta2** | **Actin, aortic smooth muscle** | **cytoskeletal protein (PC00085)** |
| **ENSMUSG00000059430** | **Actg2** | **Actin, gamma-enteric smooth muscle** | **cytoskeletal protein (PC00085)** |
| ENSMUSG00000003033 | Ap1m1 | AP-1 complex subunit mu-1 | membrane traffic protein (PC00150) |
| ENSMUSG00000026280 | Atg4b | Autophagy Related 4B Cysteine Peptidase | protein modifying enzyme (PC00260) |
| ENSMUSG00000040298 | Btbd16 | BTB/POZ domain-containing protein 16 | unclassified |
| ENSMUSG00000073077 | Cfap47 | Cilia and flagella-associated protein 47 | structural protein (PC00211) |
| ENSMUSG00000033658 | Ddx19b | RNA helicase | RNA metabolism protein (PC00031) |
| ENSMUSG00000058454 | Dhcr7 | 7-dehydrocholesterol reductase | metabolite interconversion enzyme (PC00262) |
| ENSMUSG00000016349 | Eef1a2 | Elongation factor 1-alpha 2 | translational protein (PC00263) |
| ENSMUSG00000069495 | Epc2 | Enhancer of polycomb homolog 2 | chromatin/chromatin-binding, or -regulatory protein (PC00077) |
| ENSMUSG00000022040 | Ephx2 | Epoxide Hydrolase 2 | metabolite interconversion enzyme (PC00262) |
| ENSMUSG00000061099 | Gapdhs | Glyceraldehyde-3-Phosphate Dehydrogenase | metabolite interconversion enzyme (PC00262) |
| ENSMUSG00000049649 | Gpr3 | G-protein coupled receptor 3 | transmembrane signal receptor (PC00197) |
| ENSMUSG00000046675 | Lyset | Lysosomal Enzyme Trafficking Factor | unclassified |
| ENSMUSG00000007480 | Mc5r | Melanocortin receptor 5 | transmembrane signal receptor (PC00197) |
| ENSMUSG00000025271 | Pfkfb1 | 6-phosphofructo-2-kinase/fructose-2,6-bisphosphatase 1 | metabolite interconversion enzyme (PC00262) |
| ENSMUSG00000000632 | Sez6 | Seizure protein 6 | unclassified |
| ENSMUSG00000021892 | Sh3bp5 | SH3 domain-binding protein 5 | scaffold/adaptor protein (PC00226) |
| ENSMUSG00000014606 | Slc25a11 | Mitochondrial 2-oxoglutarate/malate carrier protein | transporter (PC00227) |
| ENSMUSG00000067377 | Tspan6 | Tetraspanin 6 | scaffold/adaptor protein (PC00226) |
| **ENSMUSG00000067702** | **Tuba3a** | **Tubulin alpha-3A chain** | **cytoskeletal protein (PC00085)** |
| **ENSMUSG00000067338** | **Tuba3b** | **Tubulin alpha-3B chain** | **cytoskeletal protein (PC00085)** |
| **ENSMUSG00000058672** | **Tubb2a** | **Tubulin beta-2A chain** | **cytoskeletal protein (PC00085)** |
| **ENSMUSG00000045136** | **Tubb2b** | **Tubulin beta-2B chain** | **cytoskeletal protein (PC00085)** |
| **ENSMUSG00000062380** | **Tubb3** | **Tubulin beta-3 chain** | **cytoskeletal protein (PC00085)** |
| **ENSMUSG00000001525** | **Tubb5** | **Tubulin beta-5 chain** | **cytoskeletal protein (PC00085)** |
| ENSMUSG00000019505 | Ubb | Polyubiquitin-B | unclassified |
| ENSMUSG00000008348 | Ubc | Polyubiquitin-C | unclassified |
| ENSMUSG00000030403 | Vasp | Vasodilator-stimulated phosphoprotein | scaffold/adaptor protein (PC00226) |

Table S2: Most significant enriched mRNAs in invadosomes

**Bold**=mRNAs coding for cytoskeletal proteins

| **EnsemblID** | **Gene name** | **Protein name** | **Protein class** |
| --- | --- | --- | --- |
| ENSMUSG00000029580 | Ahcy | Adenosylhomocysteinase | metabolite interconversion enzyme |
| ENSMUSG00000026341 | Chmp4b | Charged multivesicular body protein 4b | membrane traffic protein |
| ENSMUSG00000027597 | Cnn3 | Calponin-3 | unclassified |
| ENSMUSG00000025428 | Dars | Aspartate--tRNA ligase | translational protein |
| ENSMUSG00000006273 | Dhx9 | ATP-dependent RNA helicase A | RNA metabolism protein |
| ENSMUSG00000114133 | Dlst | Dihydrolipoyllysine-residue succinyltransferase component of 2-oxoglutarate dehydrogenase complex | metabolite interconversion enzyme |
| ENSMUSG00000015733 | Eef1a1 | Elongation factor 1-alpha 1 | translational protein |
| ENSMUSG00000006699 | Eef1a2 | Elongation factor 1-alpha 2 | translational protein |
| **ENSMUSG00000038467** | **Eif3e** | **Eukaryotic translation initiation factor 3 subunit E** | **translational protein** |
| ENSMUSG00000004665 | Fxr1 | Fragile X mental retardation syndrome-related protein 1 | translational protein |
| ENSMUSG00000053931 | Hadhb | Trifunctional enzyme subunit beta | metabolite interconversion enzyme |
| ENSMUSG00000074643 | Hnrnpc | Heterogeneous nuclear ribonucleoproteins C1/C2 | RNA metabolism protein |
| ENSMUSG00000026356 | Hspa5 | Endoplasmic reticulum chaperone BiP | chaperone |
| **ENSMUSG00000042699** | **Igf2bp2** | **Insulin-like growth factor 2 mRNA-binding protein 2** | **RNA metabolism protein** |
| ENSMUSG00000004789 | Matr3 | Matrin-3 | RNA metabolism protein |
| ENSMUSG00000037742 | Mcm2 | Serine protease inhibitor A3K | protein-binding activity modulator |
| ENSMUSG00000016349 | Mvp | V-type proton ATPase 16 kDa proteolipid subunit | scaffold/adaptator protein |
| ENSMUSG00000020929 | Myof | Myoferlin | membrane traffic protein |
| ENSMUSG00000030738 | Ndufs1 | NADH-ubiquinone oxidoreductase 75 kDa subunit | metabolite interconversion enzyme |
| ENSMUSG00000022336 | Nedd4l | E3 ubiquitin-protein ligase NEDD4-like |  |
| ENSMUSG00000022337 | Phb2 | Prohibitin-2 | unclassified |
| ENSMUSG00000027680 | Ppp3ca | Serine/threonine-protein phosphatase 2B catalytic subunit alpha isoform | protein modifying enzyme |
| ENSMUSG00000026103 | Prdx2 | Peroxiredoxin-2 | metabolite interconversion enzyme |
| ENSMUSG00000026879 | Psmd13 | 26S proteasome non-ATPase regulatory subunit 13 | protein modifying enzyme |
| ENSMUSG00000059447 | Rab11a | Ras-related protein Rab-11A | protein-binding activity modulator |
| ENSMUSG00000060373 | Rack1 | Receptor of activated protein C kinase 1 | unclassified |
| ENSMUSG00000015165 | Rars | Arginine--tRNA ligase | translational protein |
| ENSMUSG00000026864 | Rbmxl1 | RNA binding motif protein |  |
| ENSMUSG00000033581 | Rnh1 | Ribonuclease H1 | RNA metabolism protein |
| ENSMUSG00000037236 | Rpl28 | 60S ribosomal protein L28 | translational protein |
| ENSMUSG00000002870 | Snx3 | Sorting nexin-3 | unclassified |
| ENSMUSG00000052105 | Spata5 | ATPase family protein 2 homolog | transporter |
| ENSMUSG00000030681 | Syncrip | Heterogeneous nuclear ribonucleoprotein Q | RNA metabolism protein |
| **ENSMUSG00000048612** | **Tkt** | **Transketolase** | **metabolite interconversion enzyme** |
| ENSMUSG00000026234 | Tpm1 | Tropomyosin alpha-1 chain | cytoskeleton protein |
| ENSMUSG00000025968 | Ugdh | UDP-glucose 6-dehydrogenase | metabolite interconversion enzyme |
| ENSMUSG00000024589 | Vdac2 | Voltage-dependent anion-selective channel protein 2 | transporter |

Table S3: Common enriched mRNAs and proteins in invadosomes

**Bold**= downregulated proteins by sieIF3E and found to be enriched in invadosomes (mRNA and protein level)

| **EnsemblID** | **Gene name** | **Protein name** | **Protein class** |
| --- | --- | --- | --- |
| ENSMUSG00000027187 | Cat | Catalase | metabolite interconversion enzyme |
| ENSMUSG00000032097 | Ddx6 | Probable ATP-dependent RNA helicase DDX6 | RNA metabolism protein |
| **ENSMUSG00000038467** | **Eif3e** | **Eukaryotic translation initiation factor 3 subunit E** | **translational protein** |
| **ENSMUSG00000042699** | **Igf2bp2** | **Insulin-like growth factor 2 mRNA-binding protein 2** | **RNA metabolism protein** |
| ENSMUSG00000032383 | Ppib | Peptidyl-prolyl cis-trans isomerase B | chaperone |
| **ENSMUSG00000048612** | **Tkt** | **Transketolase** | **metabolite interconversion enzyme** |
| ENSMUSG00000041763 | Tpp2 | Tripeptidyl-peptidase 2 | protein modifying enzyme |

Table S4: Enriched mRNAs in invadosomes deregulated in proteome from NIH-3T3 Src cells treated with sieIF3E

**Bold**= downregulated proteins by sieIF3E and found to be enriched in invadosomes (mRNA and protein level)

| **PTK ASSAY** | | | | | | | |
| --- | --- | --- | --- | --- | --- | --- | --- |
| **SRC VS WT** | | | | **SRC + PP2 VS SRC** | | | |
| **Kinase Name** | **Kinase Family** | **Mean Specificity Score** | **Median Kinase Statistic** | **Kinase Name** | **Kinase Family** | **Mean Specificity Score** | **Median Kinase Statistic** |
| LTK | Alk | 0.6 | 18.7 | CTK | Csk | 0.82 | -1.95 |
| Tyro3/Sky | Axl | 1.58 | 20.26 | EphB1 | Eph | 1.19 | -1.7 |
| EphA1 | Eph | 0.58 | 20.03 | EphA5 | Eph | 1.05 | -1.65 |
| **FAK2** | Fak | 0.54 | 17.97 | EphA10 | Eph | 0.85 | -1.59 |
| IGF1R | InsR | 0.65 | 19.2 | EphB2 | Eph | 0.82 | -1.52 |
| **JAK1~b** | JakA | 0.93 | 24.86 | **FAK2** | Fak | 1.48 | -1.92 |
| JAK2 | JakA | 0.57 | 18.12 | FAK1 | Fak | 1.1 | -1.84 |
| **Met** | Met | 1.02 | 20.22 | FGFR1 | FGFR | 1.43 | -2.89 |
| Ron | Met | 0.76 | 21.31 | FGFR4 | FGFR | 0.57 | -2.13 |
| ROR1 | Ror | 1.35 | 33.37 | **JAK1~b** | JakA | 1.06 | -1.64 |
| RYK | Ryk | 1.01 | 25.08 | **Met** | Met | 1.22 | -1.97 |
| **MEKK6/MAP3K6** | STE | 1.18 | 29.83 | PDGFR[alpha] | PDGFR | 0.94 | -1.54 |
| **ASK/MAP3K5** | STE | 1.18 | 29.83 | Brk | Src | 1.06 | -1.57 |
| MAP2K7 | STE | 0.76 | 21.89 | **MEKK6/MAP3K6** | STE | 0.98 | -1.61 |
| **MKK3/MAP2K3** | STE | 0.59 | 19.86 | **ASK/MAP3K5** | STE | 0.98 | -1.61 |
| **MKK6/MAP2K6** | STE | 0.59 | 19.86 | **SEK1/MAP2K4** | STE | 1.25 | -1.69 |
| MEK1/MAP2K1 | STE | 0.57 | 19.49 | MAP2K7 | STE | 1.1 | -1.6 |
| **SEK1/MAP2K4** | STE | 0.56 | 19.32 | **MKK3/MAP2K3** | STE | 1.05 | -1.61 |
| Syk | Syk | 1.21 | 18.57 | **MKK6/MAP2K6** | STE | 1.05 | -1.61 |
| ZAP70 | Syk | 0.96 | 18.47 | FLT4 | VEGFR | 0.81 | -1.46 |

Table S5: Pamgene analysis PTK assay

**Bold** = common deregulated kinases

| **STK ASSAY** | | | | | | | |
| --- | --- | --- | --- | --- | --- | --- | --- |
| **SRC VS WT** | | | | **SRC + PP2 VS SRC** | | | |
| **Kinase Name** | **Kinase Family** | **Mean Specificity Score** | **Median Kinase Statistic** | **Kinase Name** | **Kinase Family** | **Mean Specificity Score** | **Median Kinase Statistic** |
| **ADCK3** | ABC1 | 1.67 | 2.37 | **ADCK3** | ABC1 | 1.38 | -1.08 |
| CDK2 | CDK | 1.36 | 1.94 | AlphaK1 | Alpha | 0.96 | -1.11 |
| CDC2/CDK1 | CDK | 1.32 | 1.83 | CDK10 | CDK | 0.73 | -0.99 |
| CDK7 | CDK | 1.2 | 1.86 | **ROCK1** | DMPK | 1.65 | -1.26 |
| CDK4 | CDK | 0.78 | 1.8 | **ROCK2** | DMPK | 1.17 | -1.1 |
| CDK11 | CDK | 0.76 | 1.9 | **HIPK1** | DYRK | 1.82 | -1.44 |
| CDKL2 | CDKL | 0.81 | 2.24 | IKK[beta] | IKK | 1.68 | -1.43 |
| **ROCK1** | DMPK | 2.06 | 3.02 | PKC[zeta] | PKC | 1.91 | -1.15 |
| **ROCK2** | DMPK | 1.15 | 2.15 | PKC[beta] | PKC | 1.53 | -1.07 |
| **HIPK1** | DYRK | 0.93 | 2.44 | **PKC[iota]** | PKC | 1.5 | -1.01 |
| p38[delta] | MAPK | 1.81 | 2.06 | PKC[epsilon] | PKC | 1.43 | -0.94 |
| JNK1 | MAPK | 1.78 | 1.91 | PKC[eta] | PKC | 1.28 | -0.99 |
| JNK3 | MAPK | 1.76 | 1.91 | PKC[theta] | PKC | 1.2 | -0.91 |
| p38[beta] | MAPK | 1.22 | 2.14 | PKC[delta] | PKC | 1.07 | -0.92 |
| p38[gamma] | MAPK | 1.12 | 1.94 | RAF1 | RAF | 1.87 | -1.17 |
| JNK2 | MAPK | 1.06 | 1.79 | **MSK1** | RSK | 1.65 | -1.17 |
| MAPK14 | MAPK | 0.99 | 1.75 | **p70S6K** | RSK | 1.21 | -0.93 |
| **PKC[iota]** | PKC | 1.53 | 2.1 | RSK1/p90RSK | RSK | 1.11 | -1.07 |
| **MSK1** | RSK | 0.85 | 1.97 | RSKL1 | RSKL | 1.53 | -1.37 |
| **p70S6K** | RSK | 0.79 | 1.72 | HGK/ZC1 | STE20 | 0.97 | -1.08 |

Table S6: Pamgene analysis STK assay

**Bold** = common deregulated kinases

| **Target gene** | **siRNA ID** | **Provider** |
| --- | --- | --- |
| eIF3d human | s16499 | Ambion |
| eIF3d human | s16500 | Ambion |
| eIF3d mouse | S79900 | Ambion |
| eIF3d mouse | s79901 | Ambion |
| eIF3e human | s7485 | Ambion |
| eIF3e human | s7483 | Ambion |
| eIF3e mouse | s68369 | Ambion |
| eIF3e mouse | s68370 | Ambion |
| eIF3h human | s16509 | Ambion |
| eIF3h human | s16507 | Ambion |
| eIF3h mouse | s86276 | Ambion |
| eIF3h mouse | s86274 | Ambion |
| eIF3k mouse | s92041 | Ambion |
| eIF3k mouse | S92039 | Ambion |
| IGF2BP2 mouse | s115360 | Ambion |
| IGF2BP2 mouse | s115361 | Ambion |
| TKT mouse | 241972 | Sigma |
| TKT mouse | 241973 | Sigma |
| v-Src human | s13413 | Ambion |
| v-Src human | s13414 | Ambion |

Table S7: Lists of siRNAs used for the study

| **Antibodies** | **Source** | **Identifier** | **Application** |
| --- | --- | --- | --- |
| IgG1 Isotype Control from murine myeloma | Sigma | M5284 | IP |
| Mouse monoclonal anti-DDK | ThermoFisher | TA50011 | IF |
| Mouse monoclonal anti-eIF4E | Santa Cruz | sc-9976 | IP |
| Mouse monoclonal anti-GFP | Santa Cruz | sc-9996 | WB |
| Mouse monoclonal anti-Igf2bp2 | Santa Cruz | sc-377014 | WB/IF |
| Mouse monoclonal anti-Puromycin | Millipore | MABE343 | WB |
| Rabbit monoclonal anti-eIF3H | Cell signaling | 3413S | WB |
| Rabbit monoclonal anti-eIF4E | Cell signaling | 2067 | WB |
| Rabbit monoclonal anti-eIF4G | Cell signaling | 8701 | WB |
| Rabbit monoclonal anti-phospho-AKT(T308) | Cell signaling | 2965 | WB |
| Rabbit monoclonal anti-phospho-S6K1(T389) | Cell signaling | 9234 | WB |
| Rabbit monoclonal anti-Src | Cell signaling | 2109 | IHC |
| Rabbit polyclonal anti-4E-BP1 | Cell signaling | 9452 | WB |
| Rabbit polyclonal anti-AKT | Cell signaling | 9272 | WB |
| Rabbit polyclonal anti-eIF3A | Cell signaling | 2538 | WB |
| Rabbit polyclonal anti-eIF3B | Novus Biologicals | NBP238597 | WB |
| Rabbit polyclonal anti-eIF3C | AB Clonal | a7022 | WB |
| Rabbit polyclonal anti-eIF3D | AB Clonal | a5947 | WB/IHC |
| Rabbit polyclonal anti-eIF3E | Fortis Life Science | A302-985A | WB/IHC |
| Rabbit polyclonal anti-eIF3H | Novus Biologicals | NBP184870 | IHC |
| Rabbit polyclonal anti-eIF3I | Sigma | HPA029940 | WB |
| Rabbit polyclonal anti-eIF3K | Abcam | ab85968 | WB |
| Rabbit polyclonal anti-eIF3L | AB Clonal | A9972 | WB |
| Rabbit polyclonal anti-phospho-4E-BP1(T37/46) | Cell signaling | 9459 | WB |
| Rabbit polyclonal anti-phospho-AKT(S473) | Cell signaling | 9271 | WB |
| Rabbit polyclonal anti-phospho-Src (T419) | Cell signaling | 2101 | WB |
| Rabbit polyclonal anti-S6K1 | Cell signaling | 9202 | WB |
| Rabbit polyclonal anti-Src | Cell signaling | 2108 | WB |
| Rabbit polyclonal anti-TKT | Sigma | HPA029480 | WB |
| Rabbit polyclonal anti-TKS5 | Santa Cruz | sc-30122 | IF |

Table S8: List of antibodies used for this study

| **SequenceName** | **Sequence** |
| --- | --- |
| Mm-Igf2bp2-01 | CTGTGGGGCGACTCCCTGAGGGTATCTTACACTCGGACCTCGTCGACATGCATT |
| Mm-Igf2bp2-02 | CCTGATCTTGCGTTGTGCAGTCTGGCTTACACTCGGACCTCGTCGACATGCATT |
| Mm-Igf2bp2-03 | GGCACGATAACTTCTGCACTTGTCAAGTTCTTACACTCGGACCTCGTCGACATGCATT |
| Mm-Igf2bp2-04 | CCAGCGGTCGACGAGGGGACTCGGATTTACACTCGGACCTCGTCGACATGCATT |
| Mm-Igf2bp2-05 | ACATCTGGACCTTCTGCTGGAGCAATTTACACTCGGACCTCGTCGACATGCATT |
| Mm-Igf2bp2-06 | CTTACAGTCTCCTGCTCTGGGTAGGAGTTTACACTCGGACCTCGTCGACATGCATT |
| Mm-Igf2bp2-07 | TCTCTCGGGGTTATAAATGCTCAAATCCTGTTACACTCGGACCTCGTCGACATGCATT |
| Mm-Igf2bp2-08 | AGAATCATGCGGCATGCTTCAGATGTCCTTACACTCGGACCTCGTCGACATGCATT |
| Mm-Igf2bp2-09 | GATGTTCTTTATGGTCAAGCCCTCCTTTCTTACACTCGGACCTCGTCGACATGCATT |
| Mm-Igf2bp2-10 | CTTGTTCCCGGGCACGATGAGGGGGTTTACACTCGGACCTCGTCGACATGCATT |
| Mm-Igf2bp2-11 | AGGAGAGCTCACCTCTTCATCGGGGATTACACTCGGACCTCGTCGACATGCATT |
| Mm-Igf2bp2-12 | GTTTCTGTATCTGTGTTGACTTGCTCCACGTTACACTCGGACCTCGTCGACATGCATT |
| Mm-Igf2bp2-13 | GAATCTGGATTCTTCTGCTCCTTAGCTTTTTTACACTCGGACCTCGTCGACATGCATT |
| Mm-Igf2bp2-14 | GAGACTGAGTAGTCAACTTCCATGATTTTCCCTTACACTCGGACCTCGTCGACATGCATT |
| Mm-Igf2bp2-15 | CGGATGGCCCAGTTCTGGTCGGGGTATTACACTCGGACCTCGTCGACATGCATT |
| Mm-Igf2bp2-16 | CGAAGAGCTGCCGGAGGTCGTCGGCGTTACACTCGGACCTCGTCGACATGCATT |
| Mm-Igf2bp2-17 | ACGGCGGGGCTCAGGTTCCCAATGTATTACACTCGGACCTCGTCGACATGCATT |
| Mm-Igf2bp2-18 | CCTCATTCTCGTCTGGCGTTTGGTCACTTACACTCGGACCTCGTCGACATGCATT |
| Mm-Igf2bp2-19 | GTTCACGGTTTTCCCGCCCTTGCCAATTACACTCGGACCTCGTCGACATGCATT |
| Mm-Igf2bp2-20 | GAAGTTTTCTTCCTTCAGTTTCCCAAAGATCCTTACACTCGGACCTCGTCGACATGCATT |
| Mm-Igf2bp2-21 | CCCTGAGCCTTAAACTGGGCTTCAGGTTACACTCGGACCTCGTCGACATGCATT |
| Mm-Igf2bp2-22 | GACCAGTGATGATGACCATCCTCTCACTTACACTCGGACCTCGTCGACATGCATT |
| Mm-Igf2bp2-23 | TGATGTGGGAATGGGCCGAAATGGTGATGAGTTACACTCGGACCTCGTCGACATGCATT |
| Mm-Igf2bp2-24 | TAGCCCTGGGATCAGATTGGCTTGTTGGTTACACTCGGACCTCGTCGACATGCATT |
| Mm-Tkt-01 | CTTCGTGGTACTTGGCTGACAGCCAGGTTACACTCGGACCTCGTCGACATGCATT |
| Mm-Tkt-02 | ACCCGGATGCTGATCTTATCTTTCTTTAGATTACACTCGGACCTCGTCGACATGCATT |
| Mm-Tkt-03 | TTCTCTGGGCGGCTGGTCCGGATGAATTACACTCGGACCTCGTCGACATGCATT |
| Mm-Tkt-04 | AAATGCCCTTTGTGTTGGCTGCTAACTCTTACACTCGGACCTCGTCGACATGCATT |
| Mm-Tkt-05 | GATGGCGGCCATGCGAATCTGGTCGATTACACTCGGACCTCGTCGACATGCATT |
| Mm-Tkt-06 | CAGAAGGGCACTGTCCGGTCACGTGTGTTACACTCGGACCTCGTCGACATGCATT |
| Mm-Tkt-07 | GGAATTCTTGGTGTCTCCATCCAGGGCTTACACTCGGACCTCGTCGACATGCATT |
| Mm-Tkt-08 | GCAACGAGGTTGTCCAGCTTGTAAATTCCAGTTACACTCGGACCTCGTCGACATGCATT |
| Mm-Tkt-09 | CAGGGAGCCAGTGGCCACATCGGTGATTACACTCGGACCTCGTCGACATGCATT |
| Mm-Tkt-10 | GCTTGTTTCGGGACAGGATGCCCGTCTTACACTCGGACCTCGTCGACATGCATT |
| Mm-Tkt-11 | AGTCAGAGCTGATCTTCCTCAGGTTCAGTTACACTCGGACCTCGTCGACATGCATT |
| Mm-Tkt-12 | GGCCTCGGGTAGGAAGCCAGCTTCAGTTACACTCGGACCTCGTCGACATGCATT |
| Mm-Tkt-13 | CAGGCGATTGGCTGTGTCCTTCAGGGTTACACTCGGACCTCGTCGACATGCATT |
| Mm-Tkt-14 | CTAGCCCTTGGTGACAAGGCCTTTCACTTACACTCGGACCTCGTCGACATGCATT |
| Mm-Tkt-15 | CTTGCACAATGGCGTCCTTGTCAATACCTTACACTCGGACCTCGTCGACATGCATT |
| Mm-Tkt-16 | AGTGACCGTCACTCCAGGTTCACCCATTACACTCGGACCTCGTCGACATGCATT |
| Mm-Tkt-17 | GCTCGGGCAGAGTCTAGGATGAGTTTCCTTACACTCGGACCTCGTCGACATGCATT |
| Mm-Tkt-18 | TCCAGGGGCTTGATAGTGAAGGGATCCATTACACTCGGACCTCGTCGACATGCATT |
| Mm-Tkt-19 | CAGCAGCCAAGGCCTCATGCAGAGTTTTACACTCGGACCTCGTCGACATGCATT |
| Mm-Tkt-20 | CTTGCTCTTCAGGACCACCTTGGCTTTTACACTCGGACCTCGTCGACATGCATT |
| Mm-Tkt-21 | CCGACCTGGAAATCCTCATTGTTGCTATAATTACACTCGGACCTCGTCGACATGCATT |
| Mm-Tkt-22 | CTGCCTTCTCTGTTGCAACTCCATCGTTACACTCGGACCTCGTCGACATGCATT |
| Mm-Tkt-23 | GGAGCCACAGAGGTTGATGTTGCTCTCATTACACTCGGACCTCGTCGACATGCATT |
| Mm-Tkt-24 | GGGTGCTCCTTTTTGAAGAGCTCCGAGAAGTTACACTCGGACCTCGTCGACATGCATT |
| FLAP-Y | AATGCATGTCGACGAGGTCCGAGTGTAA |

Table S9: 5’to 3’ sequence sof the probes used for smFISH.
